# Nucleoside analogue labelling to study the cell cycle of pancreatic cancer xenografts in the chicken embryo model

**DOI:** 10.1101/2025.09.09.675135

**Authors:** Robin Colenbier, Denisa Lodoaba Jurescu, Jean-Pierre Timmermans, Johannes Bogers

## Abstract

The chorioallantoic membrane (CAM) model is an underutilised alternative model for the research into tumour xenografts. We demonstrate that in the pancreatic cancer cell lines BxPC-3 and AsPC-1, nucleoside labelling with 5-ethynyl-2’-deoxyuridine (EdU) can be multiplexed with other human cell cycle markers, (e.g., cyclin B1 and Ki-67), particularly when combined with digital image analysis, allowing human versus chicken cell classification. Furthermore, starting from the embryonic day of development 14 (ED14), we observe embryonic cells exhibiting substantial apparent cytoplasmic accumulation of synthetic nucleosides. This was observed for (F-ara-)EdU as well as for the halogenated analogues BrdU and IdU. Characterisation of these cells suggests that they are non-proliferating and possess granules and can be found in the embryonic liver of (non-)grafted embryos, as well as in the CAM itself and within xenograft sections. Importantly, these cells are unlikely to represent activated professional antigen presenting cells as they lack MHC II expression, nor do they express CD41/CD61, ruling out thrombocyte origin. Together, our findings demonstrate that the CAM model constitutes a valuable platform for studying tumour cell cycle dynamics *in vivo.* However, nucleoside labelling applications beyond ED14 may be confounded by a previously unidentified phenomenon, namely cytoplasmic nucleoside accumulation in certain chicken cells.

**Graphical abstract:** 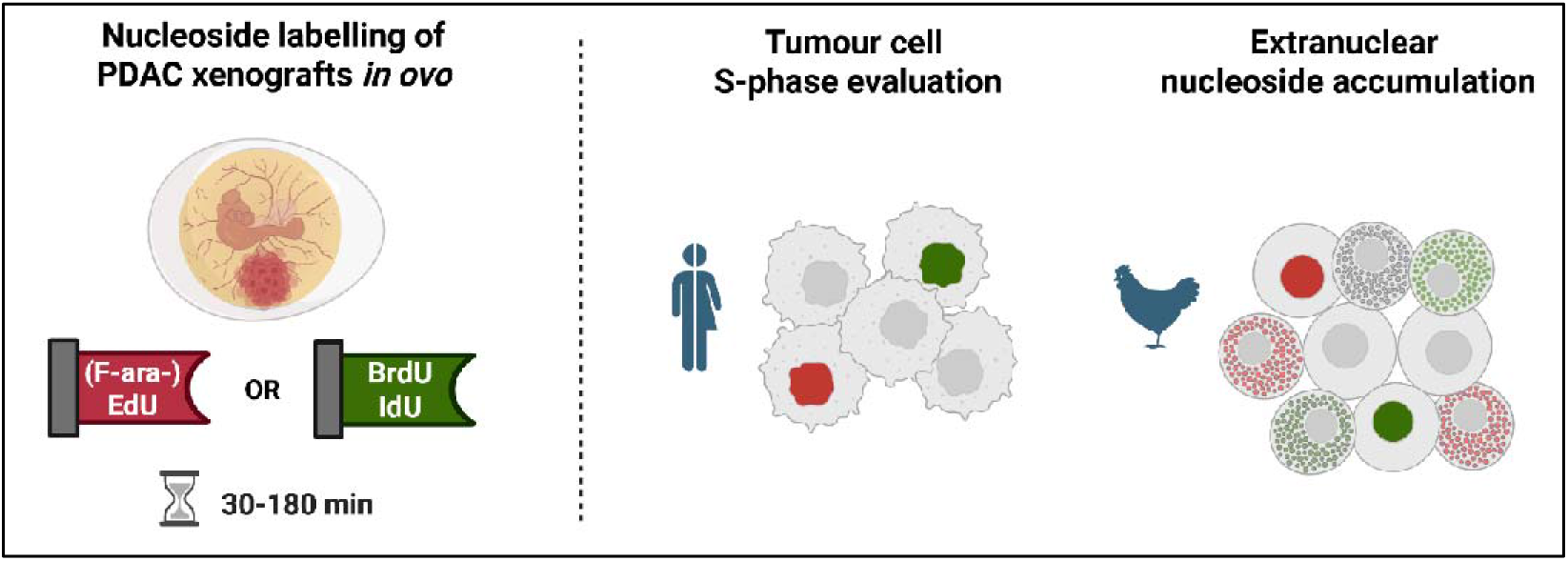

**Summary Statement:** This paper illustrates methods for nucleoside labelling in chorioallantoic membrane tumour xenografts and highlights a potential novel biological phenomenon: cytoplasmic nucleoside accumulation in granule-containing chicken cells.

## Introduction

As an alternative *in vivo* model, the use of the chorioallantoic membrane (CAM) model in cancer research is steadily gaining popularity. This is demonstrated by the growing number of original research articles employing the CAM model to evaluate cancer treatments or explore tumour biology. Despite its many advantages (e.g., low-cost, rapid turnover) the CAM assay remains underutilised in cancer research, particularly for assessing cellular proliferation and cell cycle dynamics in tumour xenografts. Furthermore, there is a scarcity of recent high-quality scientific literature regarding the use of commercial PDAC cell lines in the CAM model.(Fahmy *et al*., 2016; Costanza *et al*., 2019; Harper *et al*., 2021) Consequently, the routine application of the CAM to study tumour growth and therapy responses has not yet been widely adopted. The CAM model offers a complex biological environment in which tumour cells can proliferate in the presence of a well-vascularised extracellular matrix, connective tissue cells, and finally a (developing) immune system. One important limitation of the xenograft CAM model is the inherent sensitivity of the developing embryo to manipulations or cytotoxic treatments. As the majority of the embryo is still in full development, treatments that interfere with normal physiological processes or actively proliferating embryonic cells, could lead to decreased tumour viability and growth or may affect the performed assays at a functional level.

In this study, we demonstrate the feasibility of labelling actively dividing xenograft S-phase cells in the CAM using the halogenated thymidine analogues 5-bromo-2’-deoxyuridine (BrdU), 5-iodo-2’-deoxyuridine (IdU) as well as 5-ethynyl-2’-deoxyuridine (EdU). As both are considered to be gold standard methods for investigating the S-phase *in vitro*, their successful application in the CAM xenograft model may encourage broader use of this alternative animal model. Although already described in the context of xenografts in mice for several cancer types, these analogues are often administered intraperitoneally, which in turn leads to extended labelling durations of 3-6 hours.(Endaya *et al*., 2016; He *et al*., 2018; Chen *et al*., 2025) This impedes accurate quantification of cell cycle dynamics when a high time-resolution is required.

In contrast to rodent models, the CAM model offers easy access, allowing for tumour and embryo manipulations to be performed in a matter of seconds. This facilitates cell cycle studies in combination with exposure to compounds (such as thymidine analogues) for relatively short time-windows (i.e., < 1h). As such, the xenografted CAM model can be considered as a unique *in vivo* tool that enables both (immuno-) histological and cell cycle analyses with improved temporal resolution compared to rodent models. Notably, another alternative animal model, namely the zebrafish xenograft is being used more frequently in recent years. These models are also characterised by high accessibility and rapid development, making them compatible with quick experimental turnover and relatively high-throughput experiments. In turn, these could potentially also be used to study the cell cycle in xenograft contexts. However, no cell-cycle focused studies have been reported so far.(Yao, Wang and Wang, 2020; Somasagara and Leung, 2022; Fontana and Van Doan, 2024)

As in many cancer types, pancreatic ductal adenocarcinoma (PDAC) cell lines display considerable inter-cell line heterogeneity. Therefore, an appropriate biological model must accommodate a variety of cell lines. To this end, three distinct commercial PDAC cell lines (BxPC-3, AsPC-1, PANC-1) were used to induce tumours in the CAM. We demonstrate the feasibility of combining immunofluorescence (IF) microscopy techniques with S-phase labelling through nucleoside incorporation assays on xenograft tissue sections. To aid in the analysis and interpretation of the acquired images, digital image processing techniques were employed to differentiate proliferating tumour xenograft cells from dividing embryonic cells. Utilising the CAM model to study the cell cycle of tumour cells provides a unique and underexplored area of research that may broaden our understanding of *in vivo* cancer cell cycle regulation and lead to the improved application of currently available treatment modalities or identify potential cell cycle-related vulnerabilities of cancer cells.

## Results

### 1. The CAM model can be used to reliably induce tumour formation from commercially available PDAC cell lines

One of the primary advantages of the CAM model is its scalability. Using a relatively small setup, a large number of biological replicates can be generated within days and analysed within the following weeks. Here, we illustrate that our previously published protocol, originally optimised for the BxPC-3 cell line, can be also successfully applied for the successful xenografting of other PDAC cell lines (Figure 1A,D). Although modest success in grafting efficiencies can initially be observed when adopting the CAM model, using the proposed protocol, our lab routinely achieves tumour engraftment rates exceeding 90%, limited embryo death rates at embryonic day 10 (ED10) (12%, 10% and 9 % for the AsPC-1, BxPC-3 and PANC-1 cell lines respectively) and overall embryo survival rates of approximately 85%. Following tumour formation, the evaluated PDAC cell lines appear to retain their respective *in vitro* morphology. BxPC-3 cells exhibit a relatively well-differentiated growth morphology *in ovo,* consistent with earlier observations.(Peulen *et al*., 2013; Colenbier *et al*., 2024) These cells form clearly delineated tumour nodules embedded within the extracellular matrix, often merging into elongated, intercalated strands (Figure 1B). AsPC-1 cells also form discrete nodules, but the presence of tubular structures is less evident. In contrast, the poorly differentiated PANC-1 cells appear to be loosely interconnected and are only occasionally found as strand-like organisations, often lacking clear separation from the surrounding tumour microenvironment (Figure 1E). The nuclei of PANC-1 xenograft cells are polymorphic, closely reflecting their *in vitro* appearance (Figure 1E,F). Distinguishing tumour cells from embryonic chicken cells in the CAM model can be particularly challenging in poorly differentiated cell lines, as will be discussed further.

**Figure 1.**
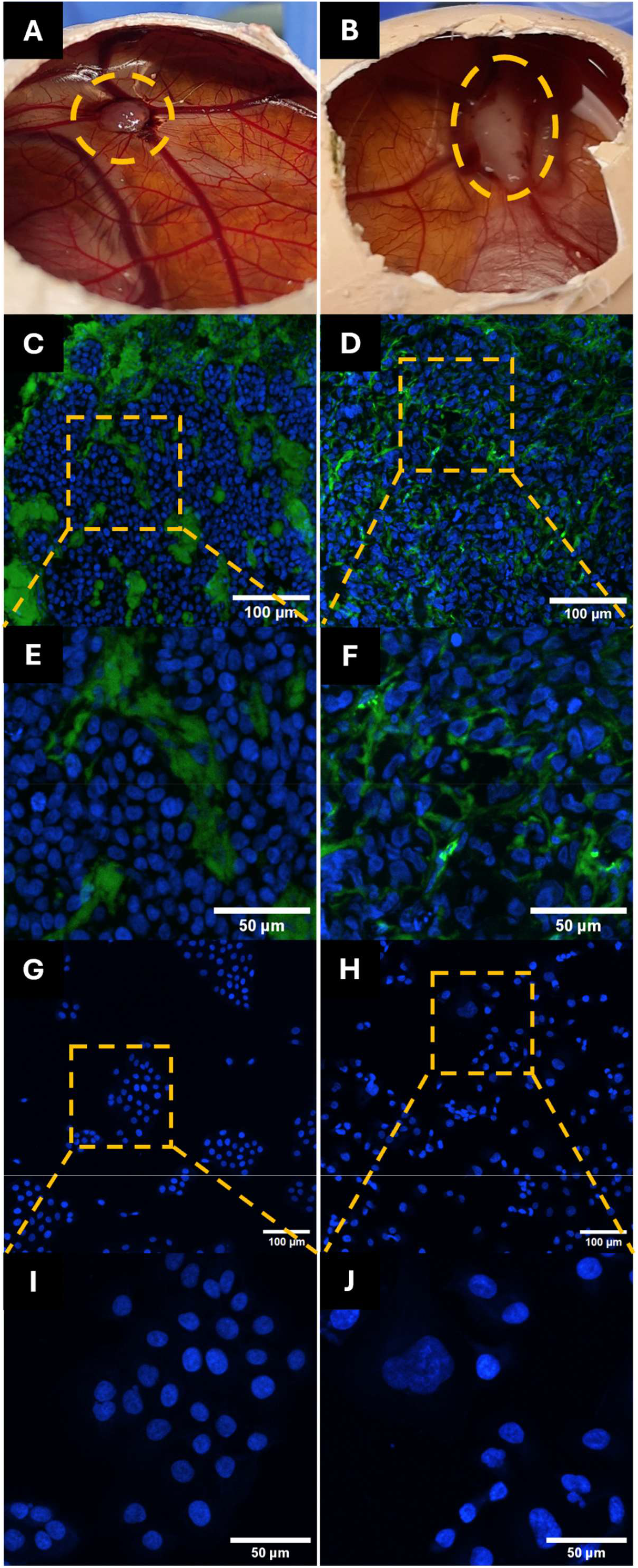
Morphological characteristics of BxPC-3 and PANC-1 cell lines *in vitro* and *in ovo.* Photographic evidence of BxPC-3 **(A)** and PANC-1 **(B)** tumour xenografts *in situ* at ED14 indicated by the yellow dashed lines. BxPC-3 xenografted cells form relatively well-defined, elongated tumour noduli **(C)** that do not appear to infiltrate into the surrounding stroma which is stained with secondary anti-chicken IgY antibodies **(green)**. In contrast, the less well-differentiated PANC-1 cell line displays a less organised and constrained growth pattern *in ovo* **(D)**. The cell-line specific growth patterns and nuclear morphological characteristics appear to be retained when comparing *in ovo **(C,E)*** and *in vitro **(G,I)*** microscopic images for the BxPC-3 cell line. *In vitro,* BxPC-3 cells appear in clearly separated groups on the overview microscopic image **(G)** and possess relatively homogenous nuclear shape and sizes (I). The morphological similarities between *in ovo **(D,F)*** versus *in vitro **(H,J)*** also appear to bre present for the PANC-1 cell line microscopic images. Here, cells appear to be grouped more randomly throughout the overview microscopic image **(H)** and PANC-1 nuclei are much more heterogenous in size as well as shape **(J)**. Tissue sections **(C-F)** are stained with secondary anti-chicken IgY antibodies **(green)**, all nuclei are stained with Hoechst 33342 **(blue)**. *In ovo* confocal microscopic images were acquired through a 20x objective; *in vitro c*onfocal microscopic images through a 40x objective.

### 2. The CAM model facilitates nucleoside incorporation assays in xenografted tumour cells

Unlike other *in vivo* models, tumour cells in the CAM model can very easily be exposed to (therapeutic) compounds due to direct access to the xenograft and its vascular supply via the (extra-)embryonic membranes. This can be exploited to label dividing xenografted tumour cells: dispensing nucleoside-containing solutions onto the CAM takes mere seconds, in turn facilitating short-term investigations. Once the solution is applied onto the CAM, compounds can be absorbed into the blood stream of the embryo and subsequently be perfused through the tumour tissue. Historically, halogenated thymidine analogues such as BrdU and IdU have been employed routinely to investigate the S-phase *in vitro* as well as *in vivo*, the latter predominantly being rodent models. More recently, halogenated analogues have been supplanted by (copper-based) click-labelling techniques which rely on exposure to compounds such as EdU and F-ara-EdU. These newer analogues offer superior specificity and multiplexing compatibility compared to the halogenated analogues. Despite their advantages and widespread use in *in vitro* experiments, all the mentioned thymidine analogues have, somewhat surprisingly, not been adopted yet in the setting of the CAM model to investigate the cell cycle of tumour xenograft cells.

#### 2.1 BrdU labelling enables S-phase detection in xenografted PDAC tumour cells via immunohisto-fluorescence

As a proof-of-concept, BxPC-3 tumour-bearing embryos at various stages of development (ED10-ED14) initially were exposed to BrdU (up to 800 µmol/kg for labelling durations up to two hours) prior to sacrifice. No clear signs of acute embryo toxicity were observed and intense S-phase labelling could be achieved in as little as 30 minutes of exposure to BrdU (Figure 2).

**Figure 2.**
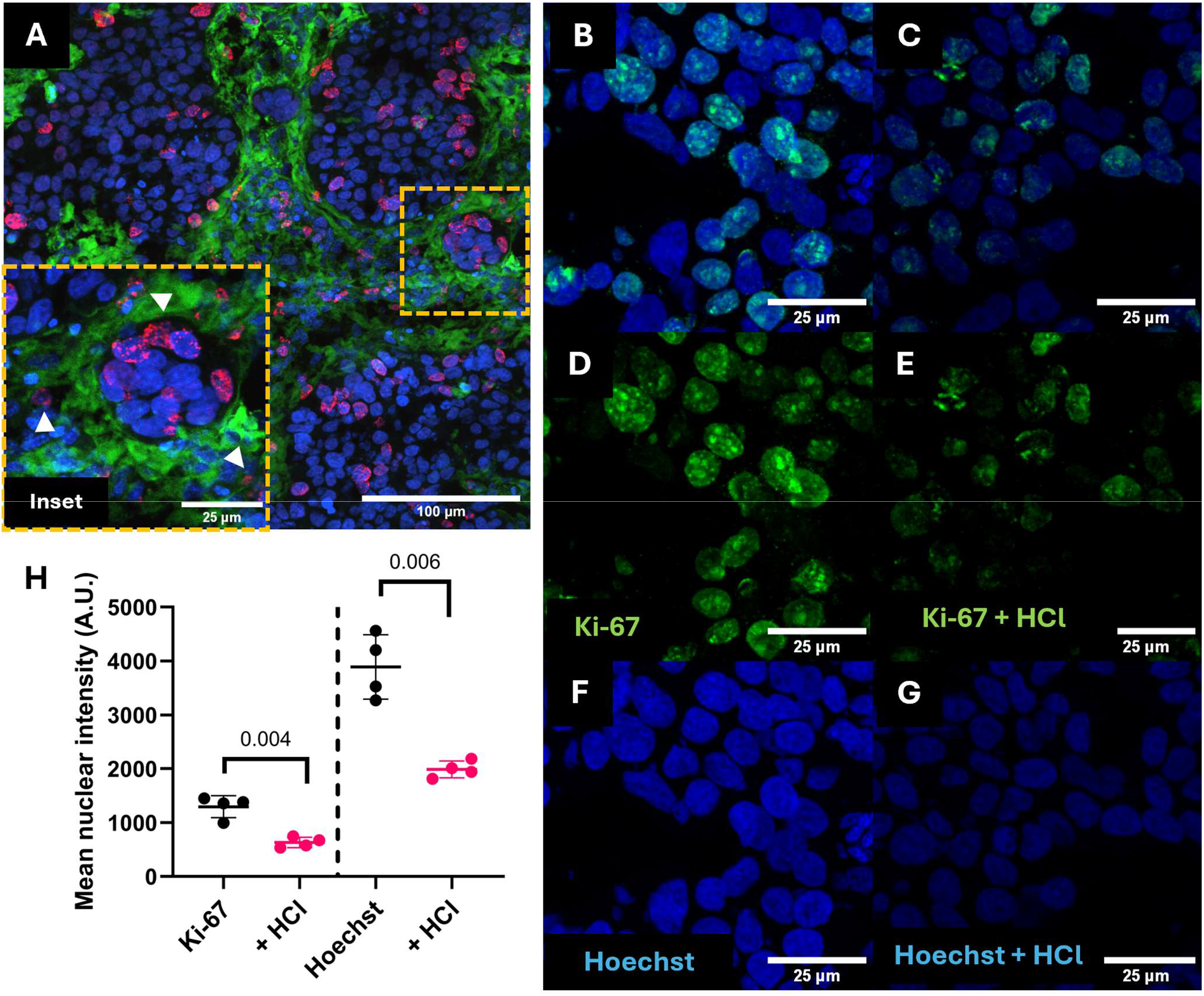
BrdU-based nucleoside labelling of a BxPC-3 xenograft. **A.** Confocal microscopic image of a BxPC-3 xenograft cryosection labelled *in ovo* for 30 minutes with BrdU (800 µmol/kg) prior to sacrifice at ED13 and followed by 60 minutes HCl-based denaturation to uncover the BrdU-epitopes with subsequent IF detection for BrdU **(red)**, and counterstaining with secondary anti-chicken IgY antibodies **(green)** and Hoechst 33342 **(blue)**. Throughout the tissue section, tumour cell S-phases can be detected through the nuclear localisation of the BrdU signal in BxPC-3 cells, which typically gives rise to a peripheral nuclear staining pattern which is indicative of incomplete dsDNA denaturation. As nucleoside incorporation assays are not species-specific, BrdU-positive suspected chicken embryonic cells nuclei **(arrowheads)** can occasionally be observed throughout the tissue sections. Pre-treatment of BxPC-3 tissue sections with HCl, required to uncover the BrdU epitopes, leads to significant reductions in subsequent nuclear staining or immunofluorescence labelling. An untreated BxPC-3 cryosection **(B)** displays intense nucleolar Ki-67 labelling along with a fairly homogenous nuclear background intensity **(D)** in addition to an intense Hoechst 33342 labelling **(F)**. HCl-treatment of tissue sections prior to subsequent labelling leads to decreased Ki-67 as well as Hoechst 33342 intensities **(C,E,G)**. In addition, the nucleolar localisation of Ki-67 IF appears to be absent in HCl-treated tissue sections **(E)**. Analysis of the mean nuclear intensities for Ki-67 IF and Hoechst 33342 staining in n = 4 technical replicates reveals that both staining procedures are significantly affected by HCl-denaturation. As a result, interpretation of signal intensities and localisation following denaturing pretreatments should be avoided. Significant differences were assessed via two-sided Welch’s t-test. ** = 0.05 > p < 0.001. Error bars represent SD. Images acquired through a 40x objective.

The BrdU signal typically localised to the nuclear periphery (Figure 2A), a characteristic pattern resulting from the obligate double-stranded DNA denaturation step (commonly using 2M HCl) required for the detection of halogenated analogues via immunodetection. As described by Pierzyńska-Mach et al.(Pierzyńska-Mach *et al*., 2016), this pattern may signify incomplete dsDNA denaturation. Nevertheless, harsher or prolonged denaturation can increase background staining and interfere with subsequent immunodetection of other markers. In extreme cases, it can lead to near-complete destruction of dsDNA helices, resulting in diminished or absent nuclear staining with standard dyes such as DAPI or Hoechst 33342 (Figure 2 F-H). Thus, standard BrdU detection protocols are incompatible with subsequent reliable IF detection of other epitopes in xenografted tumour cells. To illustrate the latter, anti-human Ki-67 IF labeling was performed following a brief BrdU detection protocol (2M HCl denaturation for 30 minutes). It is clear that two characteristic aspects of Ki-67 staining, namely the homogenous nuclear signal and an intense nucleolar pattern in proliferating, non-mitotic cells are both lost following the HCl denaturation, thereby rendering its interpretation unreliable (Figure 2 B-E, H). Performing the denaturation and BrdU detection after IF labelling will also affect fluorescence intensities due to the destruction of fluorophores.

Importantly, nuclear incorporation of BrdU is observed in both dividing human and chicken embryonic cells (Figure 2B, arrowheads), confirming its broad application potential and high sensitivity to also detect S-phase cells but highlights the absence of species-specificity. Hence, strategies are needed to avoid misidentification of embryonic cells as proliferating tumour cells. These may include human-specific IF labelling (see section 2.2) or nuclear feature-based classification via imaging processing software. Illustrating the latter, human PDAC tumour nuclei are generally larger than chicken embryonic nuclei, which is especially overt in AsPC-1 xenografts (Figure 3) while many chicken nuclei in the CAM also demonstrate intense and discrete nucleolar staining patterns with nuclear dyes.

**Figure 3.**
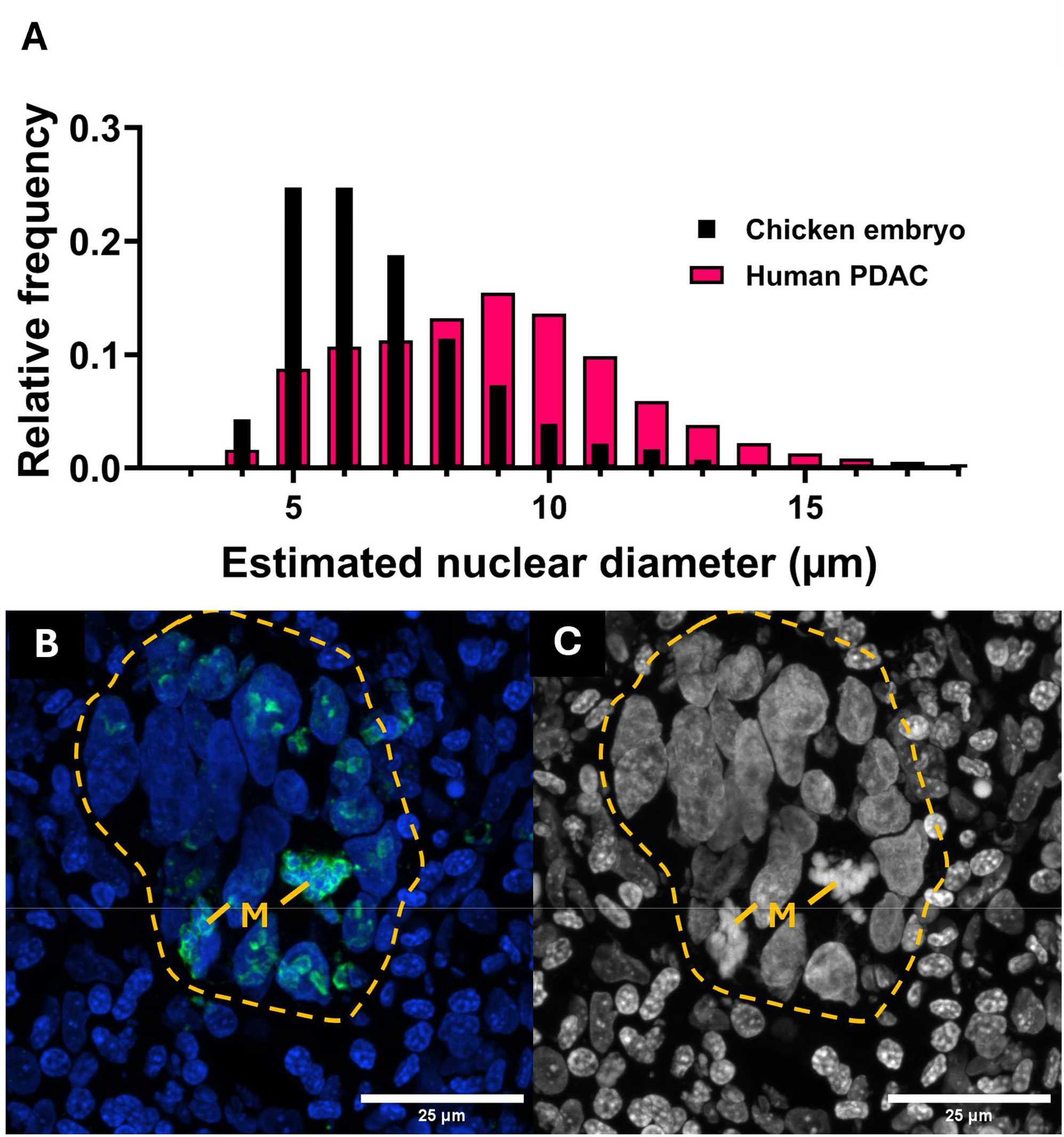
Comparison of nuclear size and morphology between human PDAC cells and chicken embryonic cells. **A.** Estimated nuclear size distribution of human PDAC nuclei versus CAM-resident chicken embryonic nuclei. In general, the chicken embryonic nuclei found within the CAM possess smaller nuclear diameters compared to human PDAC cells. This size difference can be used to aid the generation of microscopy-based species-specific classification models. However, due to the considerable overlap, additional variables will need to be taken into account for accurate predictions. Nuclear diameters were calculated through the analysis of microscopic imaging data of n = 3250 chicken embryonic CAM nuclei, n = 17692 BxPC-3 nuclei and n = 13653 AsPC-1 nuclei found in CAM xenograft tissue sections. Values for BxPC-3 and AsPC-1 cell lines were pooled to allow species-wise nuclear size comparison. **B.** Illustration of nuclear sizes and morphology of AsPC-1 cells (indicated by dashed yellow lines) surrounded by chicken embryonic nuclei within a CAM xenograft section. IF for human Ki-67 (green) illustrates the human origin of these nuclei. Mitotic nuclei (M) are identified by the perichromosomal presence of Ki-67 and should not be mistaken for dying cells. C. Gray-scale representation of the nuclear channel, highlighting the distinct nucleolar staining pattern of the chicken embryonic nuclei which may be incorporated as an additional variable in species-classification models. Nuclei are stained with Hoechst 33342. Ki-67 intensities were adjusted to maximise the visualisation of its localisation in mitotic cells. Images were acquired through a 60x objective

Nevertheless, reliance solely on nuclear features may be accompanied by various pitfalls. For example, the nuclear staining intensities obtained via Hoechst and DAPI dyes are negatively affected in a non-linear fashion by the amount of nucleoside incorporation due to the distortion of the dsDNA helix.(Rausch *et al*., 2021) This phenomenon occurs even in the absence of denaturation steps (e.g., as required for BrdU detection). Therefore, the use of these nuclear, dye-dependent intensity measurements for species classification becomes challenging in the context of nucleoside labelling, especially when relying on BrdU-based labelling strategies. Lastly, (cell-line dependent) cellular heterogeneity can further impede effortless classification, especially in dense multicellular environments, such as xenografted tumours. Consequently, additional IF labelling is required to segregate proliferating chicken and human cells. Replacing BrdU with EdU enables reliable detection of other (cellular) markers in parallel as the detection of EdU does not rely on denaturation treatments for its detection.

#### 2.2 The application of 5-ethynyl-2’-deoxyuridine (EdU) allows specific S-phase labelling and multiplexing with immunofluorescent labelling of cell cycle markers in xenografted tumour cells

Unlike halogenated thymidine analogues, detection of EdU within tissue sections does not require dsDNA denaturation steps, making it highly compatible with IF staining (Figure 4). Moreover, BrdU signal intensities do not faithfully represent the quantity of incorporated analogue as its detection is heavily dependent on the applied denaturation protocol. In contrast, it is generally accepted that EdU signal intensities do closely reflect the degree of nuclear incorporation. After immunolabeling for human Ki-67, we confirmed the specificity of both markers in xenograft nuclei (Figure 4E) and the compatibility of both labelling strategies for xenograft tumour cells within the CAM. For example, Ki-67^+^ nuclei that were not in S-phase at the moment of nucleoside labelling (e.g., mitotic nuclei) clearly lacked any EdU signal (Figure 4A-C). Importantly, anti-Ki-67 IF showed retained fidelity and perichromosomal localisation in mitotic nuclei following the EdU click reaction, demonstrating the method’s suitability for studies across multiple cell cycle phases (Figure 4D). Using the proposed methodology (see section 2.3.1), approximately 95% of EdU^+^ tumour nuclei are correctly attributed a Ki-67-positive status (Figure 4E). In addition to Ki-67 detection, the EdU labelling in CAM xenografts can be combined with other cell cycle markers, such as (human) cyclin B1, enabling finer resolution of the different cell cycle stages of the cancer xenograft *in vivo* via microscopy. Cytoplasmic cyclin B1 presence can be observed in the late S-phase during the transition to- and throughout the G2 phase. The bi-phasic localisation of cyclin B1 during the cell cycle, namely cytoplasmic during S/G2 and increasingly localised to the nuclear compartment during late G2/mitosis offers extra dimensionality.(Hwang *et al*., 1995; Arooz *et al*., 2000; Frisa and Jacobberger, 2009; Karuna *et al*., 2020) Combined, these markers allow *in ovo* segregation of proliferating tumour cells into early S-phase (ES; EdU^+^CB1^cyto-^), late S-phase/early G2-phase (LS; EdU^+^CB1^cyto+^) and G2-phase (EdU^-^CB1^cyto+^) (Figure 4E-G).

**Figure 4.**
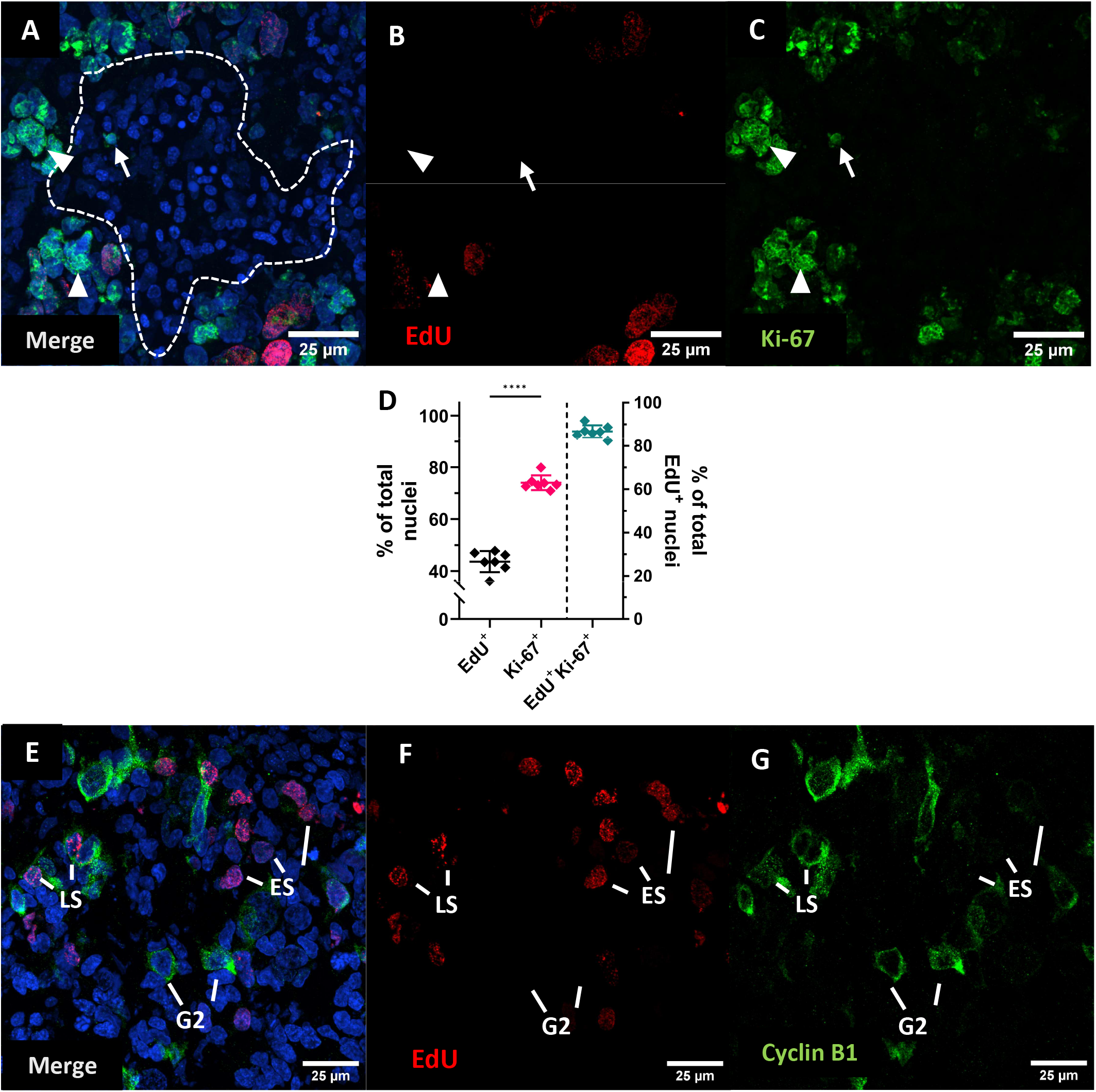
Successful combination of Ki-67 and Cyclin B1 immunofluorescence labelling with EdU detection in CAM xenografts. A-D. Cryosection from an AsPC-1 xenograft (ED15) labelled with EdU (red, 800 µmol/kg for 30 minutes) and stained for Ki-67 **(green)**. The white dashed region outlines an embryonic cell population clearly negative for human Ki-67. The Ki-67 positive nuclear fragment **(arrow)** belonging to an AsPC-1 nucleus in a different tissue plane should not be misinterpreted as a (false Ki-67^+^-positive) chicken nucleus. **Arrowheads** depict Ki-67^+^, but EdU^-^ mitotic tumour nuclei, illustrating the specificity of both staining techniques for different cell cycle stages within the described application and is confirmed by the quantification of marker positivity in tissue sections derived from n = 7 xenografts **(E)**. Plot respectively illustrates the calculated percentage of EdU^+^ (Ki-67^+/-^) over the total number of tumour nuclei (44%), Ki-67^+^ tumour nuclei over the total number of tumour nuclei (74%) and the percentage of tumour nuclei that are correctly calculated to be EdU^+^ and Ki-67^+^, respective to the total number of EdU^+^ tumour nuclei (94%). As a result, the applied techniques succeed in correctly identifying Ki-67-positivity in the vast majority of all detections. Statistical testing was performed using one-way ANOVA with Tukey’s post-hoc test: **** = p < 0.0001; error bars represent SD. **F-I**. Cryosection of an AsPC-1 xenograft (ED13) labelled with EdU (**red**, 800 µmol/kg for 30 minutes) and IF-labelled for cyclin B1 **(green)**, illustrating the compatibility of the *in ovo* combination of EdU labelling with different cell cycle markers. Here, the combination of nuclear EdU incorporation and the cytoplasmic localisation of cyclin B1, in the absence of nuclear cyclin B1, allows segregation of xenograft cells into (early) S-phase **ES**, late S-phase (**LS**) and G2 phase (**G2**). Images were acquired through a 60x objective.

#### 2.3 Considerations for applying nucleoside labelling in the CAM model to investigate the cell cycle in xenografts

As mentioned previously, the microscopic evaluation of nucleoside labelling of xenografts clearly demonstrated the presence of BrdU^+^ or EdU^+^ chicken embryonic cells in close proximity to xenograft tumour cells. As nucleoside incorporation is not species-specific, the use of techniques to distinguish human (tumour) versus embryonic chicken cells is warranted. In essence, the evaluation of the cell cycle of xenografted tumours in the CAM needs to be combined with additional techniques (such as the addition of human-specific IF labelling) to allow accurate human versus chicken segmentation of acquired images. This way, proliferating chicken embryonic cells can be excluded from analyses.

##### 2.3.1 Application of both specific and non-specific immunofluorescence techniques facilitates tissue segmentation

In order to avoid the false inclusion of (dividing) chicken embryonic cells in analyses, several experimental strategies can be employed that cooperate with digital image processing (e.g., via the QuPath software package).(Bankhead *et al*., 2017) First, distinguishing between both species can already be achieved to a large extent in a relatively simple manner via the use of anti-chicken IgY fluorophore-conjugated antibodies, which are routinely mainly used as secondary antibodies. Although we prefer to allocate the green light spectrum (e.g. FITC) for this application, anti-IgY labelling can of course be performed with other fluorophores, such as Cy3 (Supplementary Figure S1). In doing so, chicken embryonic tissue components are intensely stained (Supplementary Figure S2). Importantly, the IgY present within the embryonic tissues is not expressed by the embryonic cells themselves at this stage of development, but is derived from the egg yolk.(Carlander, Wilhelmson and Larsson, 2003) The IgY pool residing in the yolk has been transferred from the mother hen to provide passive immunity. Subsequent IgY transfer from yolk to embryonic tissues takes place continuously throughout the embryonic development, allowing its detection in the embryo.(Dias da Silva and Tambourgi, 2010; Ulmer-Franco, 2012)

Second, nuclear features can be extracted and used to aid in the generation of a classifier model using built-in algorithms included in the QuPath software. This can already result in modestly accurate tissue classification for relatively well-differentiated cell types, such as BxPC-3. Species distinction can be further facilitated by combining anti-chicken IgY IF (Figure 5C,H) with additional anti-human IF, such as anti-human Ki-67 (Figure 5D), in comparison to classification based upon solely nuclear characteristics (Figure 5B). Alternatively, one can opt to apply a binary classifier based upon anti-IgY IF by manually setting a defined intensity threshold based upon visual inspection of the pixel intensity distribution in the “anti-chicken” channel (Figure 5E). However, the latter option may result in higher numbers of false-negative tumour cell detection.

**Figure 5.**
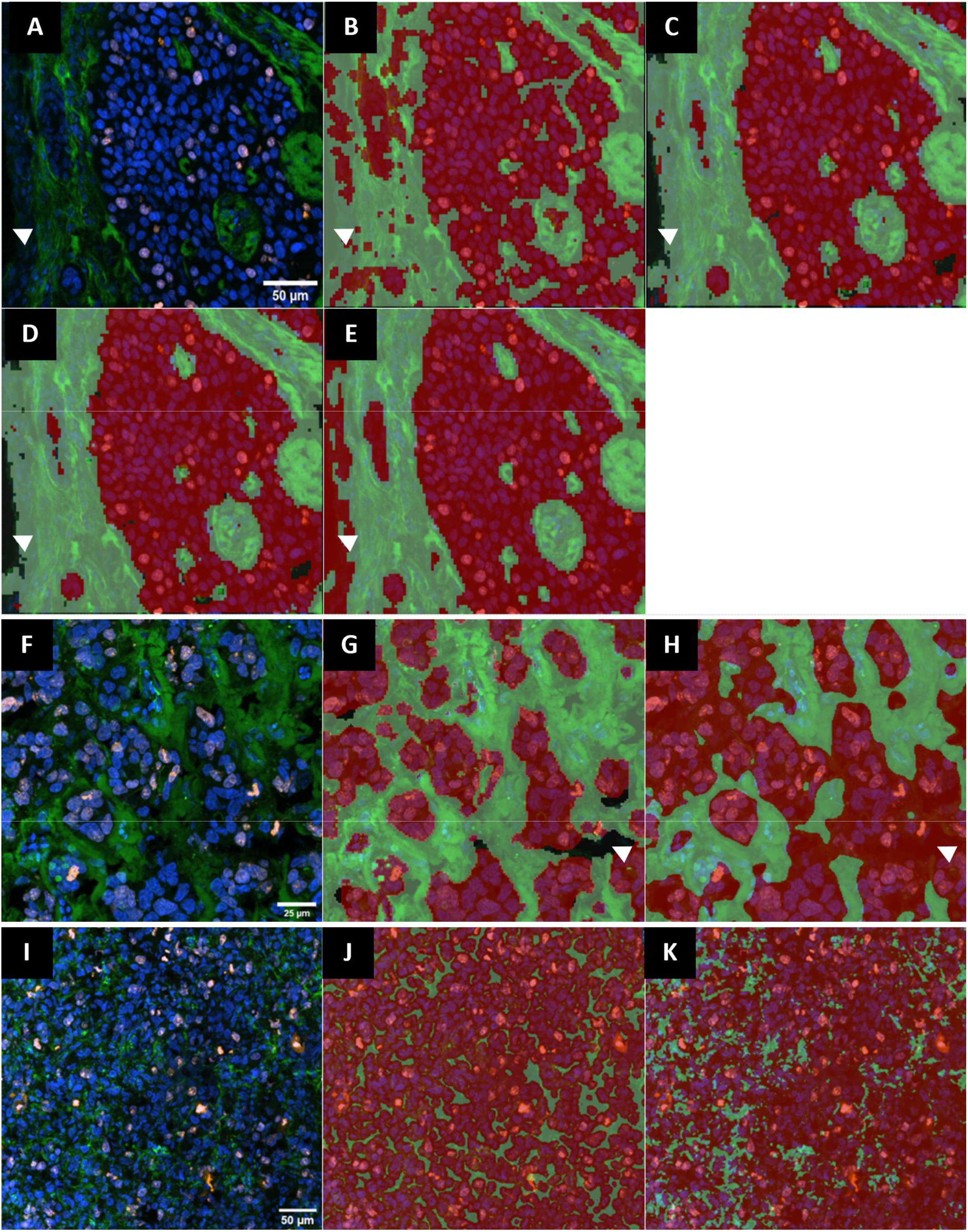
Comparison of different pixel classification strategies to discriminate human PDAC cells (red) from chicken tissue (green) within the CAM. **A.** Raw confocal image of a BxPC-3 xenograft cryosection at 20x magnification. **B.** Trained classifier output based solely upon nuclear channel image data. **C.** Trained classifier output taking into account nuclear channel image data as well as anti-IgY IF labelling shows improved segregation of tumour cells, chicken stroma and empty (non-tissue) areas **(arrowheads). D.** Addition of anti-Ki-67 IF channel image data does not considerably alter the classifier output, suggesting anti-IgY IF is sufficient to allow species-wise segmentation in well-differentiated PDAC xenografts. **E.** The application of a binary thresholder, via manual assignment of a fixed intensity threshold based on visual inspection and quantification of the signal intensities in the anti-IgY IF channel allows adequate human versus chicken distinction but introduces bias via classifying empty (non-tissue) areas as tumour areas **(arrowheads). F.** Raw image of an AsPC-1 tumour cryosection at 60x magnification **G.** Trained classifier output, taking into account anti-IgY and anti-Ki-67 IF image data. **H.** Similar to BxPC-3, the application of a binary thresholder classification strategy (anti-IgY IF) allows low-level tissue segmentation but can introduce false positive tumour areas **(arrowheads)** for AsPC-1 xenografts. I. Raw image of a PANC-1 xenograft section at 20x magnification. **J.** Application of a two-channel trained classifier (anti-IgY and anti Ki-67 IF) leads to only moderately successful tissue classification, in part as a result of the poorly-defined growth pattern of PANC-1 cells. **K.** Consequently, the binary thresholding strategy is also not successful for species-wise cell segregation in PANC-1 xenografts. Trained classifiers were designed per cell line using the ANN_MLP-model, built in the QuPath software package (see Materials & Methods). Sections were stained with Hoechst 33342 **(blue)**, anti-Ki-67 **(orange),** anti-IgY **(green)**. Arrowheads indicate non-tissue areas. Classifier outcomes (**tumour = red** versus **chicken = light green**) are indicated by the coloured translucent overlay.

Human versus chicken segmentation based on the binary threshold strategy performs adequate in cases where a clear distinction between tumour and chicken tissues is already present to some extent. This is often observed in xenografts of well-to moderately-differentiated cell lines (i.e., BxPC-3, AsPC-1). However, it is only is reliable provided there is limited technical variability in staining results across replicates. Nevertheless, this strategy may also induce artifacts, as non-tissue (empty) areas can be classified as tumour (Figure 5, arrowheads). In turn, a higher extent of manual curation will be warranted, which can introduce additional observer bias and impact processing speed. In contrast, for poorly-differentiated cell lines such as PANC-1, the performance of the simple thresholder is limited (Figure 5K).

When human cells can be confidently separated from chicken tissue, by (supervised) automated or manual annotation, individual cell or nuclei detection can be performed more accurately. In this step, detection of nuclei can be performed using the built-in *“cell detection”* functionality in QuPath. This can be performed for the entire image (Figure 6B,F) or only for the tumour region(s) of interest (Figure 6 C,G). Once nuclei have been segmented, targets can be quantified in a human-specific manner. Not applying species-specific masks or classification strategies may lead to the inclusion of a significant amount of chicken nuclei in subsequent analyses and introduce bias in experimental outcomes as illustrated by the species-wise nuclei quantification using the various strategies (Figure 6 D,H).

**Figure 6.**
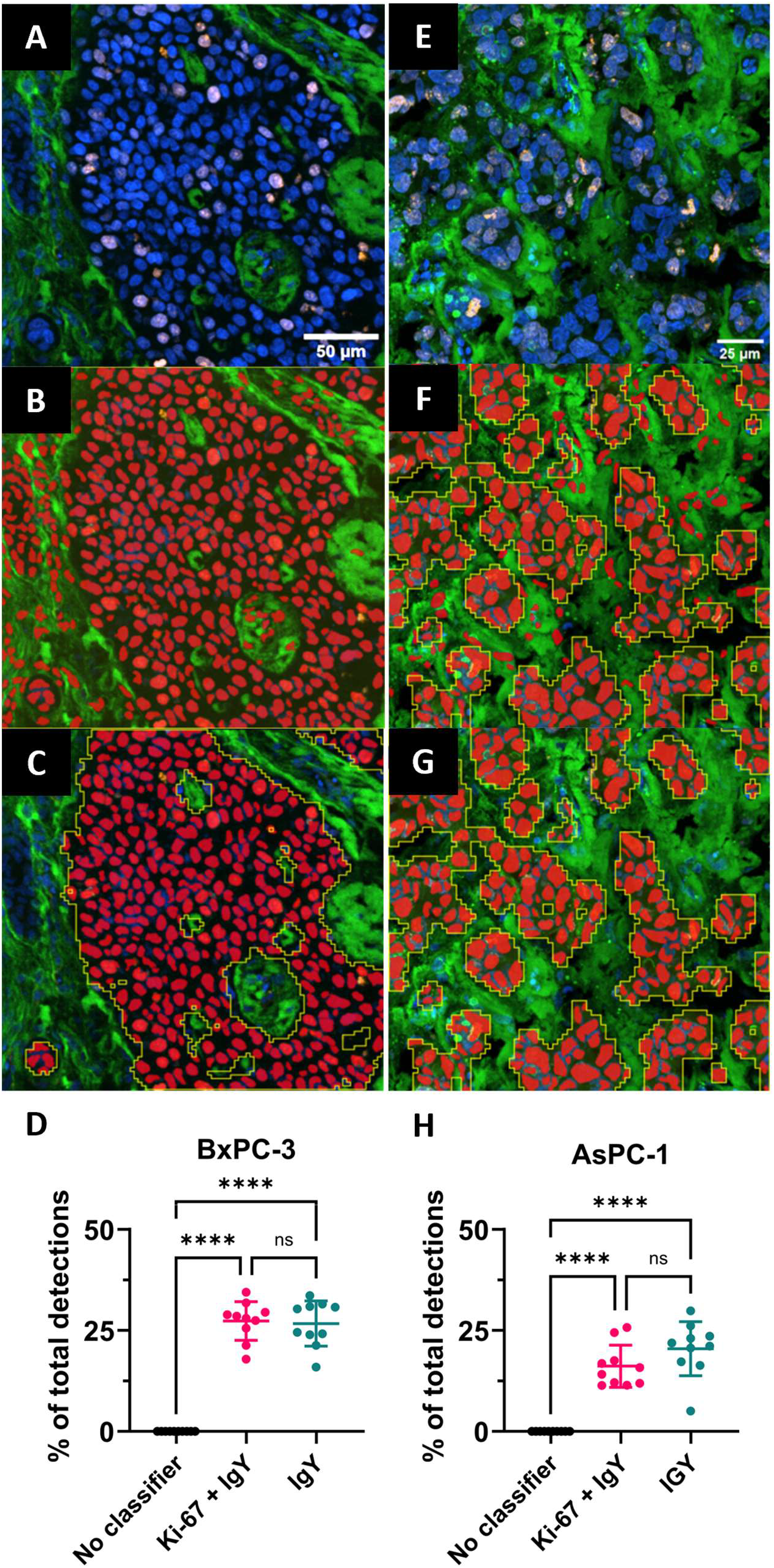
Combination of a trained two-channel classifier (anti-IgY and nuclear channel) with the cell detection plugin from QuPath facilitates distinction between human and chicken embryonic nuclei. **A.** Identical BxPC-3 cryosection as depicted in Figure 5A. **B.** Running the cell detection plugin without species-wise classification strategies severely overestimates the number of tumour nuclei (red) within the tissue section. **C.** Application of a trained two-channel classifier prevents chicken nuclei to be classified as human PDAC nuclei and thus increases the accuracy of subsequent quantification. **E.** Identical AsPC-1 cryosection as depicted in Figure 5F. **F.** Not applying species-wise classification strategies consistently overestimates the number of tumour nuclei within a xenograft section, which can in turn contribute to false positive nucleoside-positive detections. **G.** Application of a trained two-channel classifier allows the removal of false-positive nucleus detections. **D,H.** Comparison of two different classifier strategy outcomes, i.e., two-channel trained classifier and binary thresholder (IgY channel) for AsPC-1 and BxPC-3 tissue sections (n = 10). In both cases, the application of a species-wise classification strategy allows the identification and consequent removal of false-positive tumour nuclei detections, thereby improving fidelity of subsequent analyses. Statistical testing was performed using one way ANOVA with Tukey’s post-hoc test: **** = p < 0.0001; ns = p ≥ 0.05; error bars represent SD. Sections are labelled with anti-chicken IgY **(green)**, anti-human-Ki-67 **(orange)** and Hoechst 33342 **(blue)**. Bright red overlay signifies detected nuclei, yellow lines indicate the area correctly classified as tumour.

##### 2.3.2 Causes of variability in apparent EdU-labelling efficiency: emergence of a specific but non-nuclear EdU signal

When applying nucleoside labelling in the CAM model to investigate the cell cycle in xenografts, the labelling solution containing the analogue should be dispensed onto the CAM as far away from the xenograft as possible. The rationale is that in doing so, the majority of the analogue that will reach the tumour, will have passed through the embryonic cardiovascular system first. This way, the effects of passive diffusion through the CAM (starting from the dispensing area) on the incorporation of the nucleoside analogue in the tumour will be kept to a minimum. Embryonic liver sections of non-grafted embryos, stained for EdU confirmed that the analogues pass through the embryonic circulation illustrated by the clear presence of EdU^+^ hepatoblast or hepatocyte nuclei (Figure 7A). Surprisingly, cells with perinuclear, granular, and intracytoplasmatically localised EdU accumulation were observed within the liver parenchyma (Figure 7A-B). A clear overlap of the nucleoside signal with cytoplasmic granules can be observed (Figure 7C-D). These cells were initially only sporadically observed within embryonic liver sections at ED14 although they appeared more frequently with increasing embryonic development stage (≥ ED14). Eventually, this phenomenon was consistently observed within the surrounding stroma of post-ED14 CAM xenografts (Figure 7C-D). The presence of mitotic figures within the xenograft suggests that the absence of nuclear EdU labelling of the tumour cells was not the result of non-proliferation but could be the result of nucleoside sequestration in the chicken cells. Importantly, the cells harbouring the granular cytoplasmic nucleoside signal are cells are not apoptotic, as demonstrated by the absence of active-caspase-3 IF labelling (Figure 7E-F).

**Figure 7.**
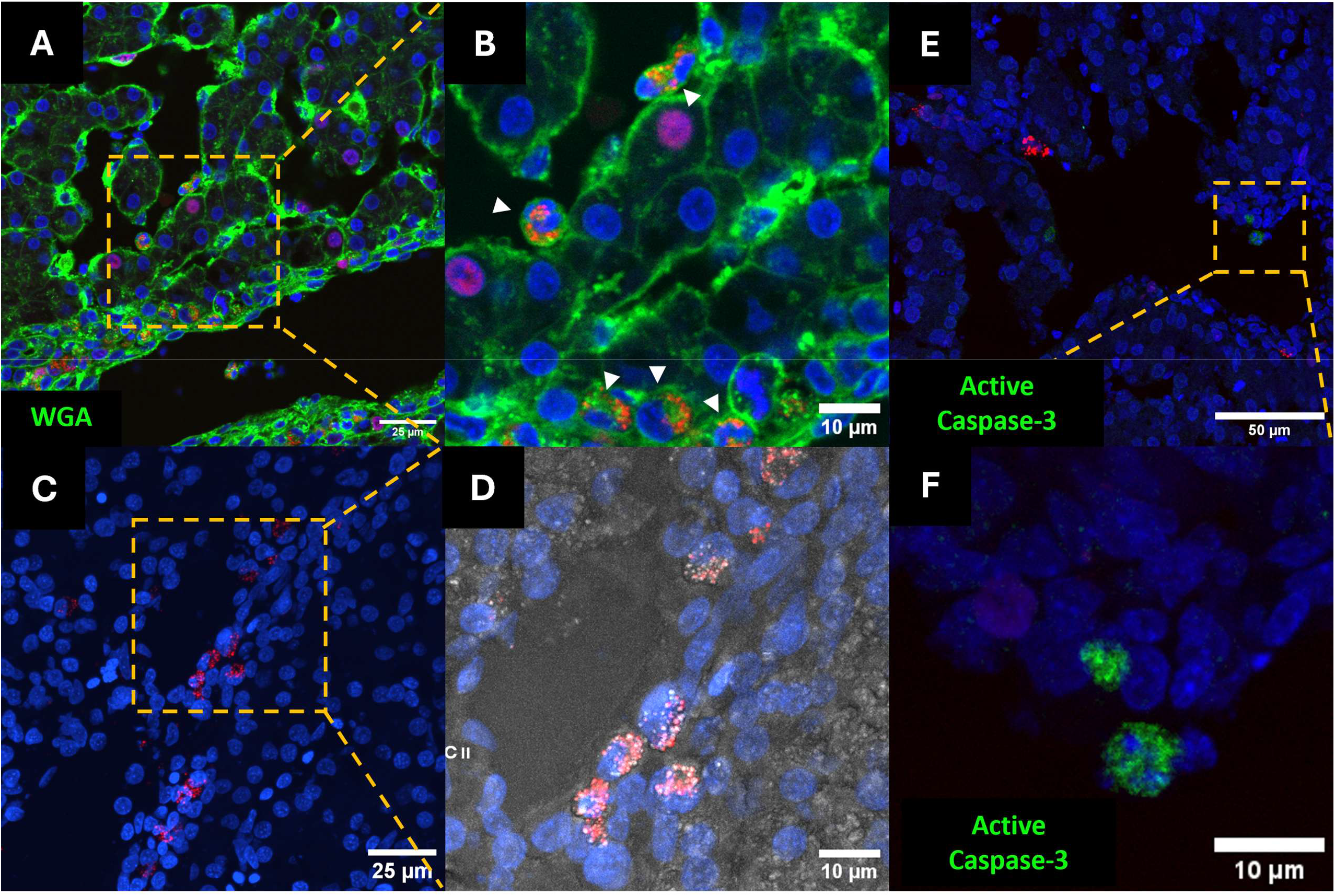
Cytoplasmic and nuclear nucleoside detection within chicken embryonic tissues. Detection of EdU (**red**, 800 µmol/kg for 30 minutes) within **A.** Embryonic liver tissue section (ED18) acquired through a 40x objective. Cell membranes are labelled with wheat-germ-agglutinin **(WGA, green)**. Nuclear nucleoside signal is confined to the liver cells while the extranuclear signal is observed in the cytoplasm of distinct cells **(arrowheads) B.** Detail view of the marked section from panel A. **C.** Embryonic liver tissue section (ED16) labelled for F-ara-EdU **(red). D.** Detail view of the marked section from panel C with brightfield imaging overlay. **E.** Chicken embryonic liver (ED16) sections containing F-ara-EdU **(red)** and stained for active caspase-3 **(green)** and subjected to the click-detection reaction confirm that the granular cells are not apoptotic. Spontaneous apoptotic cells can however be observed throughout the developing embryonic liver. **F.** Detail view of the region marked in panel E, illustrating the fragmented nucleus of the apoptotic cell. The specificity of the primary antibody was confirmed via the detection of intrinsic apoptosis in normally developing chicken embryos (see supplementary Figure S6). Nuclei are stained with Hoechst 33342 **(blue)**. Images acquired through a 40x objective.

We initially considered a generalised inflammatory response following in-lab chicken embryo incubation, tumour grafting, or the reported intrinsic toxicity of EdU to be a contributing factor.(Zhao *et al*., 2013, pp. 5-; Haskins *et al*., 2020a) Therefore, additional labelling experiments were performed on healthy, non-xenografted embryos using BrdU, IdU and F-ara-EdU which possess reduced cytotoxic potential compared to EdU.(Neef and Luedtke, 2011a; Haskins *et al*., 2020b) Following a brief denaturation protocol (30 minutes of 2 M HCl), IdU and BrdU were detected in the nuclei of embryonic liver cells (Supplementary Fig. S3). Under these conditions, the clear extranuclear BrdU or IdU signal appeared to diminish greatly and could not be detected as reliably (Supplementary Fig. S3A-B, arrows). However, IF detection of IdU and BrdU *without* prior HCl treatment revealed that the extranuclear signal was also present for the halogenated nucleoside analogues (Supplementary Fig. S3C-D). The absent detection of the extranuclear signal following denaturation suggests that the observed signal is unlikely to be the result of (ds)DNA fragments that contain the halogenated analogue. If this were the case, the denaturation would generally lead to increased labelling intensities. Moreover, this demonstrates that the occurrence of the extranuclear signal for BrdU and IdU is not induced by the denaturation protocol.

Furthermore, labelling of embryos with F-ara-EdU (650 µmol/kg for 3 hours) showed identical staining patterns to those observed for EdU and for IdU and BrdU under non-denaturing conditions (Supplementary Fig. S3-S4). Nuclear incorporation rates for F-ara-EdU during replication are reported to be much lower compared to EdU and the halogenated analogues, which was also apparent within the chicken tissue sections.(Neef and Luedtke, 2011b) Interestingly, the extranuclear accumulation of F-ara-EdU signal appeared to be present as abundantly as to that which had been observed for EdU and BrdU/IdU. This again, makes it less likely that the extranuclear signal originated from nucleoside-containing DNA, considering the slow nuclear incorporation rate of F-ara-EdU. For all evaluated analogues, clusters of cells containing the accumulated nucleosides tended to localise near larger vessel lumina within the embryonic liver (Supplementary Fig. S4). Potential confounding (bio)chemical causes for the detected cytoplasmic nucleoside accumulation were considered; IF detection of EdU with a cross-reactive anti-BrdU antibody did not lead to discernible extranuclear EdU signal (not shown). Moreover, addition of a click-detection reaction buffer on tissue sections of embryos without nucleoside labelling, or with BrdU or IdU, did not reveal the typical observed staining pattern (Supplementary Figure S5). As mentioned, the cells containing cytoplasmic nucleoside labelling exhibited a granular appearance. Thus, we sought provide further insight into the possibility that these cells represented some type of antigen-presenting cells. Therefor, immunofluorescent labelling directed to chicken MHC class II was carried out. In both liver and xenograft sections, extranuclear-EdU containing cells were found to be negative for MHC II, confirming that these cells are likely not *professional* antigen-presenting cells (Figure 8A-B). In lower vertebrates, including avians, phagocytic functions are also ascribed to thrombocytes. Existing research shows that fluorescent beads and dyes can accumulate within the cytoplasm of (avian) thrombocytes, resulting in similar staining patterns to our observations.(Carlson, Sweeny and Tokaryk, 1968; DaMatta, Seabra and de Souza, 1998; Sugimoto, Nishikawa and Sugiyama, 2023) Nevertheless, immunofluorescent labelling for the avian homolog of CD41/CD61 as a marker for thrombocytes demonstrated that the nucleoside signal accumulation did not take place within CD41^+^/CD61^+^ cells (Figure 8C-F). Furthermore, the cytoplasmic aspect visible via brightfield imaging of nucleoside-positive cells (Figure 8F) was distinct from that of the CD41^+^/CD61^+^ cells. Clearly, the cytoplasmic nucleoside signal was also not observed in (nucleated) chicken erythrocytes (Figure 8F).

**Figure 8.**
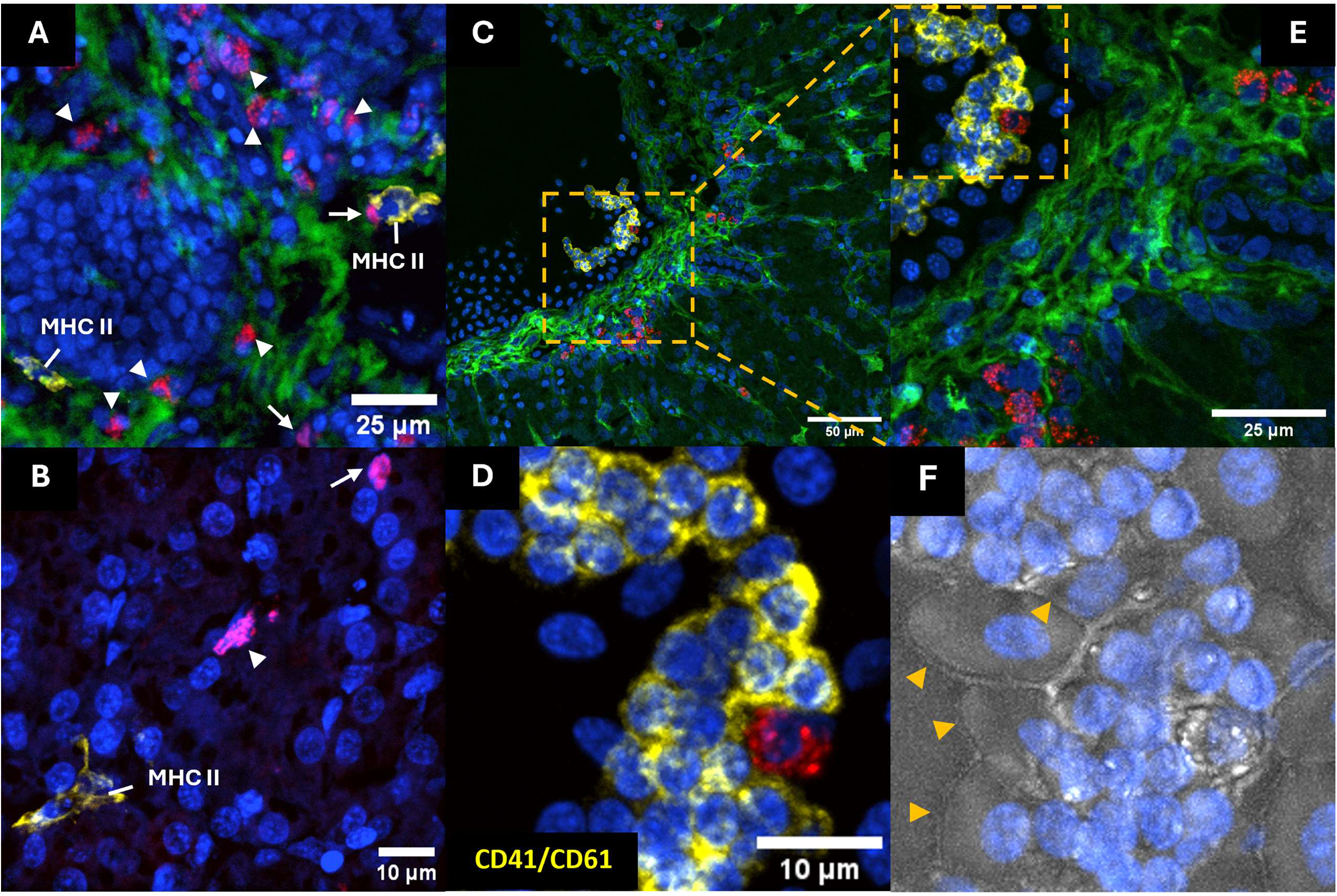
Illustration of nuclear and cytoplasmic nucleoside detection in three different tissue sections. **A.** BxPC-3 xenograft tissue section (ED18) demonstrating near-absent nuclear EdU incorporation **(arrows)** while cytoplasmic inclusions in surrounding non-tumour cells are abundantly present **(arrowheads)** these inclusions were not observed within the soma of MHC II^+^ cells **(yellow)** present in the CAM, acquired at 20x magnification. Chicken tissue is labelled with anti-IgY IF **(green)**. **B.** Cytoplasmic EdU was not detected within MHC II^+^ cells residing in the chicken embryonic liver (ED18) of non-grafted embryos, acquired at 40x magnification. **C.** Chicken embryonic liver (ED18) labelled with F-ara-EdU (red), anti-chicken CD41/CD61 **(yellow)**, and anti-chicken IgY **(green)**. **D-F.** Within the blood vessel, nucleated erythrocytes **(yellow arrowheads)** are present in close proximity to the thrombocytes and a granular, cytoplasmic F-ara-EdU containing cell. Neither erythrocytes or thrombocytes display nuclear or cytoplasmic nucleoside signal. Image acquired at 40x magnification. Nuclei are stained with Hoechst 33342 **(blue)**.

## Discussion

### Nucleoside labelling of tumour xenografts in the CAM model is feasible and should be paired with species-directed classification strategies

In this paper, we have demonstrated that our previously published protocol concerning the xenografting of the BxPC-3 cell line (Colenbier *et al*., 2024) can also be applied to the AsPC-1 and PANC-1 cell lines. Additionally, the *in vitro* growth pattern of these cell lines appears to be retained well during *in ovo* xenograft culture. Consequently, distinguishing the moderately to well-differentiated cell lines BxPC-3 and AsPC-1 from embryonic tissues is relatively straightforward. In contrast, poorly differentiated cell lines, which exhibit a high degree of cellular heterogeneity, impede the clear segregation of xenografts from CAM tissues. Furthermore, we demonstrated that BrdU, IdU and EdU enable efficient labelling of dividing xenograft nuclei within the CAM model with excellent labelling efficiency achievable in as little as 30 minutes. No obvious toxicity was observed with either compound, but the toxicity profile of EdU in particular should be taken into account as multiple reports have shown that prolonged exposure can lead to DNA damage accumulation and cytotoxicity.(Zhao *et al*., 2013; Neef and Luedtke, 2014; Haskins *et al*., 2020a; Wang *et al*., 2022) For longer duration S-phase labelling, the more recently developed analogue F-ara-EdU can be used. While F-ara-EdU also offers high multiplexing potential and appears to have very low toxicity, it exhibits much slower nuclear incorporation kinetics compared to EdU and BrdU/IdU.(Neef and Luedtke, 2011b) Therefore, F-ara-EdU is not suitable for short-term cell cycle labelling for xenograft procedures such as those described in this paper. Nevertheless, EdU retains superiority to BrdU or IdU in terms of sensitivity and compatibility with subsequent IF. The latter is particularly important given the non-species-specific incorporation of nucleosides into S-phase nuclei in the CAM model.

Next, we have presented simple yet effective techniques that aid in the analysis of tumour cell cycle in the CAM xenograft model, namely the application of anti-IgY IF staining and the use of at least one anti-human IF marker in combination with built-in classifier models in the QuPath software package. When combined, these approaches provide excellent results in terms of segregating human xenograft nuclei from chicken embryonic nuclei in AsPC-1 and BxPC-3 xenografts. This strategy is applicable to microscopic images acquired at 20x, 40x and 60x magnification. Importantly, this workflow allows imaging and processing of whole-tissue sections rather than imaging and quantifying select *representative* tissue areas – as is commonly performed. We are convinced that similar segregation strategies should be applied in any xenograft tumour model, including rodent models, to minimise observer bias in the quantification of tumour tissues. A possible limitation of this study is that examples presented were generated from the relatively well-to-moderately defined tumour xenografts derived from the BxPC-3 and AsPC-1 cell lines (as opposed to PANC-1 tissues). However, to date, there are no commercially available, nor validated primary antibodies that can specifically detect (nuclear) chicken antigens which would allow segregation between human and chicken cells within tumour xenografts. Our proposed strategies offer accessible methods that can aid overcoming this limitation, in turn increasing the application potential of the CAM model in cancer research, but they do not appear to as effective on xenograft tissue sections derived from poorly differentiated cell lines. However, we are confident that these examples sufficiently demonstrate the applicability of nucleoside labelling in CAM xenografts when it is combined with species-wise cell classification. Therefore, optimisation and fine-tuning of classifier models that perform robustly across a variety of cell lines fall outside the scope of this paper.

### A newly identified phenomenon: extranuclear/cytoplasmic nucleoside accumulation in chicken embryonic cells

In addition to the consistent labelling of S-phase cells with nucleosides within both embryonic liver sections and xenograft nuclei, we identified at least one chicken embryonic cell population that appeared to contain high amounts of extranuclear nucleosides. This phenomenon was also observed in liver sections of embryos that did not receive tumour grafting and was finally confirmed to be present for the analogues IdU, BrdU and F-ara-EdU in subsequent experiments. Critically, in all of the evaluated conditions and tissue types, the presence of the cytoplasmic nucleoside signal within one cell excludes the presence of nuclear nucleoside signal within the same cell and *vice versa*. As a result, it is very unlikely that cells demonstrating the cytoplasmic accumulation are actively proliferating at that moment, nor in the three hours following incorporation.

Following initial observations, several control staining procedures were performed to rule out staining-related technical artefacts. Non-specific reactivity due to high amounts of (reactive) free thiol groups present on proteins could result in significant interference. For example, Wiest & Kielkowski demonstrated that in biological samples, the simultaneous presence of both azide-and alkyne-groups, used in the copper-catalysed click reaction, can interact with the aforementioned thiol moieties and result in a detectable thiotriazole product.(Wiest and Kielkowski, 2024) This reaction may lead to ubiquitous background labelling in cell lysate samples when low concentrations of ascorbic acid (i.e., ≤ 1 mM) are used in the reaction buffer. However, in our tissue staining procedure, we applied an ascorbic acid concentration of 5 mM at which the thiotriazole formation is negligible.(Wiest and Kielkowski, 2024) This is also supported by the absence of ubiquitous background labelling in the surrounding tissues of the cells with cytoplasmic nucleoside signal within (F-ara-)EdU containing tissue slides. In non-nucleoside containing tissue sections, addition of the click-detection buffer did not lead to the emergence of this signal. Indeed, it would be exceptionally unlikely for thiotriazoles to be formed in the absence of azide-moieties in the tissue sections. The simultaneous presence of cells with cytoplasmic nucleoside accumulation alongside cells showing the expected nuclear incorporation in both embryonic liver and xenografted tumour nuclei further diminishes the likelihood of confounding technical issues that hamper the efficiency of the click-detection reaction.

This extranuclear accumulation was also investigated in tissues gathered from embryonic livers from both grafted and non-grafted embryos. None of these tissues showed nuclear or cytoplasmic staining, confirming the absence of potential ubiquitous background signal induction. Next, liver and xenograft sections from EdU-exposed embryos were incubated with an anti-BrdU antibody known to exhibit substantial cross-reactivity with DNA-incorporated EdU.(Liboska *et al*., 2012) In both tumour and liver tissues, the cross-reactive anti-BrdU antibody failed to reproduce the extranuclear EdU signal in any embryonic cells. Therefore, it is unlikely that the extranuclear EdU signal represents (fragmented) EdU-containing DNA. Interestingly, extranuclear presence of BrdU/IdU could only be reliably detected when tissues were not subjected to prior denaturation through 2M HCl, a prerequisite for the uncovering of DNA-incorporated halogenated analogues. This also suggests that the observed signal is not derived from BrdU/IdU-containing DNA. The fact that we were able to demonstrate reproducibility and localisation of the signal with two independent techniques, using their complementary analogues (i.e., click-detection for (F-ara-)EdU and immunofluorescence detection for BrdU/IdU) further strengthen our findings.

It is also evident that the apparent presence of cytoplasmic nucleosides in cells is not confined to the liver-resident cells, as this phenomenon was also observed in CAM-resident cells. These cells likely represent a ubiquitously distributed cell population present in both healthy and xenografted chicken embryos. Given that embryos were exposed to nucleoside labelling for no longer than three hours, it is highly unlikely that these cells are a progeny, induced by proliferation or differentiation in response to high concentrations of the potentially toxic analogues. Since cytoplasmic nucleoside-containing cells are consistently present in non-grafted embryos as well, we can conclude that the phenomenon is not triggered by xenografting or the presence of PDAC cells. Although initially observed in liver tissue sections, we cannot exclude the possibility that these cells may originate in other tissues or organs (e.g. spleen or the Bursa of Fabricius) since no other tissues have been examined so far.

As one potential explanation for the observed phenomenon, we considered the signal to be indicative of extranuclear (fragmented) DNA present in damaged or apoptotic cells. Thus, the extranuclear signal would be indicative of replicated DNA (during the labelling period) that had fragmented (e.g. via caspase-3/7-dependent cleavage). However, we have not observed nuclear fragmentation or pyknosis in cells displaying cytoplasmic nucleoside accumulation. IF labelling for active-caspase-3 confirmed the absence of apoptosis in these cells. Additionally, the extranuclear signal was never observed in cells with clear nuclear labelling. This does not exclude the possibility that cells harbouring the cytoplasmic nucleoside accumulation are situated within early apoptotic stages. Alternatively, we hypothesised that the cells containing the cytoplasmic nucleoside signal were a ubiquitous type of macrophages. However, an assessment for (activated) professional antigen-presenting cells (APC) via IFM for monomorphic chicken BLA chain (MHC class II) revealed that these cells did not carry this marker and are thus unlikely to be professional APC. On the other hand, avian thrombocytes are known to carry out phagocytic capabilities (Carlson, Sweeny and Tokaryk, 1968; DaMatta, Seabra and de Souza, 1998; Ferdous and Scott, 2023), but IFM for CD41/CD61 showed that the extranuclear accumulation did not take place in CD41/CD61-positive cells. Lastly, brightfield imaging suggested the presence of granules within the cytoplasm of cells that contained the extranuclear nucleoside signal. It is also clearly visible that the signal overlaps with cytoplasmic granules. Given the high number of extranuclear nucleoside-positive, CD41^-^/CD61^-^, granule-containing cells, these may represent heterophils; the avian functional homolog of neutrophils. Similar to mammalian biology, these are also the dominant white blood cell type in the chicken embryo.(Genovese *et al*., 2013) Additionally, the liver is a source of granulopoiesis starting from ED15 (Guedes *et al*., 2014), the same time point at which we started to observe large amounts of cells with cytoplasmic nucleoside signal. To further elucidate the identity of these cells, in-depth investigations using avian markers are required which may prove challenging as only a limited number of characterisation markers are available for the avian immune system.(Härtle *et al*., 2024) At this point, the observed granule-containing cells are likely to mainly represent heterophils though we have not been able to completely identify and characterise these cells due to the aforementioned lack of available primary antibodies directed versus avian markers of interest for these findings.

Nevertheless, these observations pose an important limitation for the use of nucleoside labelling in xenografts within the CAM model during later stages of embryonic development (i.e., beyond ED14). Due to the observed, presumed replication independent, cytoplasmic accumulation of the nucleosides in the embryonic cells, incorporation of the analogue into proliferating cells can be substantially reduced, potentially resulting in many false-negative detections of S-phase tumour cells. Assuming that these cells are part of the developing embryonic immune system, their impact on nucleoside labelling efficiency of tumour grafts should be minimal at earlier time points, when the immune system is still very immature and these cells are not present yet in substantial quantities. To preserve the sensitivity and specificity of nucleoside labelling, we therefore recommend performing this assay prior to ED14 in CAM xenograft experiments if the experimental aim is to investigate the cell cycle of the tumour cells.

While the presence of cells with observed extranuclear nucleoside accumulation presents a technical challenge for the interpretation of nucleoside labelling in CAM xenografts, these findings also raise several novel and important research questions:

1. Is this phenomenon caused by non-specific reactivity of the azide-dyes with endogenous molecules, and if so, through what (bio-)chemical mechanism?

If this represents a biological phenomenon:

2. Do these cells actively accumulate nucleosides or synthetic analogues, and if so, what triggers the phenomenon?
3. Is it limited to the chicken embryo model, or is it also present in different species?
4. Do the observed cells truly represent a single cell type, or can multiple distinct cell types exhibit similar behaviour?

Answering these questions could provide deeper insight into fundamental biological processes and the cellular handling of nucleosides and their analogues. Alternatively, they can provide crucial information on potential limitations for the use of nucleoside labelling *in vivo*.

## Conclusions

- IdU/BrdU as well as EdU and F-ara-EdU labelling can be used to label S-phase nuclei of proliferating cells in the CAM model in as little as 30 minutes, surpassing the time resolution obtained in rodent models. However, the implementation of additional species classification strategies is warranted to prevent misinterpretation of the incorporation assays.
- The superiority of EdU over halogenated analogues for short-term cell cycle labelling is retained, as evidenced by its multiplexing capacity, which is essential for accurate, species-specific digital quantification of microscopic images.
- We describe a novel phenomenon: Ubiquitous cytoplasmic accumulation of nucleoside analogue signal in non-proliferating, non-apoptotic, chicken granule-containing cells in embryonic chicken liver and CAM tissue.

## Materials and methods

### Cell lines and -culture

The used cell lines BxPC-3 (CRL-1687), AsPC-1 (CRL-1682) and PANC-1 (CRL-1469) were obtained from ATCC. Cells were maintained and expanded at 37 °C in a humidified 5% CO_2_ atmosphere in T175 culture flasks (Greiner Bio-One, Cat#660175) in Dulbecco’s modified Eagle’s medium, containing 4.5g/L glucose and supplemented with 4 mM + L-glutamine (Gibco, Cat#41965039), 1 V/V% Penicillin-Streptomycin (10,000 U/mL) (Gibco, Cat#15140122) and a final concentration of foetal bovine serum of 10 V/V% (Gibco, Cat#A5256701). All cell lines were routinely tested (every three months) for Mycoplasma contamination. For *in vitro* microscopic image acquisition, cells BxPC-3 and PANC-1 cells were seeded in a 96-well plate (Cellvis, Cat#P96-1.5P) at a density of 7500 cells per well on the day prior to microscopic imaging. On the day of imaging, cells were fixed for 15 minutes in a 4% formalin solution prior to further staining and processing.

### Embryo incubation

Fertilised (ED0) chicken eggs (*Gallus gallus domesticus*) were obtained from Belgabroed NV and set to incubate in an egg incubator (Fiem MG 140/200 Rurale) with relative humidity set point at 55% and temperature set point at 37 °C for xenografting experiments. For the positive control tissues for the apoptosis assay, control embryos were incubated at 37 °C. Front and hind legs were dissected at ED 7 through ED 7.5.

### Xenografting in the chorioallantoic membrane model

Xenograft tumours were generated according to our previously published protocol(Colenbier *et al*., 2024) with the following modifications: All grafting was performed at ED7 using a final cell suspension volume of 50 µL rather than 100 µL. The final amount of grafted cells per embryo was 4 million for each cell line. During the preparation of the grafting cell suspension, PANC-1 cells were centrifuged at 400 G for 5 minutes rather than 250 G for 7 minutes.

### Nucleoside labelling of dividing cells in the CAM model

5-Bromo-2⍰-deoxyuridine (BrdU, Sigma-Aldrich, Cat#19-160), 5-Iodo-2⍰-deoxyuridine(IdU, Sigma-Aldrich, Cat# I7125) and 5-Ethynyl-2’-deoxyuridine (EdU, Thermo Scientific Chemicals, Cat#466382500) were dissolved in dimethylsulfoxide (DMSO) to make a stock concentration of 100 mM and 50 mM for IdU/BrdU and EdU respectively. (2YS)-2Y-Deoxy-2Y-fluoro-5-ethynyluridine (F-ara-EdU, Sigma-Aldrich, Cat#T511293) was dissolved in distilled in distilled H_2_O at a stock concentration of 12.5 µM. Subsequently, stock solutions were sterile filtered through 0.20 µm pore size polytetrafluoroethylene (PTFE) filters (Sartorius, Cat#17575-K) and stored at -20 °C until further use.

Working solutions for *in ovo* labelling of the cell cycle were prepared by diluting each stock solution in sterile Dulbecco’s phosphate-buffered saline (DPBS, Cat#14190169). For example, for labelling with doses of 800 µmol/kg egg weight, a working solution of 13.33 mM (3.36 mg/mL for EdU, 4.09 mg/mL for BrdU) was prepared and warmed to 37 °C.

Prior to labelling, each egg was weighed and the amount of volume to be pipetted onto each CAM was calculated based upon the following formula: 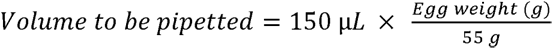

For example, an egg weighing 58.3 g would receive 159 µL of the 13.33 mM labelling solution. Then, for each egg, the appropriate amount of nucleoside-containing working solution was dispensed onto the CAM as far away from the visible tumour as possible before returning them to the egg incubator. Two researchers performed the labelling in tandem, ensuring that an experimental group of 16 embryos could be labelled in less than five minutes.

### Tissue collection and fixation

At the end of the labelling period, eggs were placed on, and covered with ice in order to halt the cell cycle and stop nucleoside incorporation. Subsequently, embryos were sacrificed via decapitation, tumours were excised from the CAM and embryo livers removed the chicken embryo. All collected tissues were immediately rinsed three times with ice-cold PBS and fixed overnight in an ice-cold 4% formalin solution (ROTI®Histofix, Cat#3105).

### Tissue (pre-)processing and cryosectioning

Following overnight fixation, tissues were extensively washed in PBS on a rotating shaker for 10 minute and a minimum of five changes of PBS. Tumours were further dissected with the aid of a stereomicroscope in order to remove excess CAM and blood clots. Next, tissues were pre-embedded overnight at 4 °C in a 30 W/V% sucrose solution in PBS containing 0.1 % sodium azide. The following day, pre-embedded tissues were embedded in NEG-50 cryo-embedding medium (Epredia, Cat#6502), equilibrated to -25 °C and eventually stored at -80 °C until further use.

7 µm thick cryosections were cut from xenograft tissues, while for embryonic liver, 10 µm thick sections were cut on pre-coated glass slides (VWR, Cat# 631-0108), dried at 37 °C for 2 hours and stored at -20 °C.

### Immunofluorescence labelling of cryosections

Cryosections were thawed, air-dried and a barrier was drawn around the tissue sections using a hydrophobic barrier pen (Sigma-Aldrich Cat#Z377821). Then, sections were rehydrated and washed in PBS for at least three washes of 10 minutes each, before being covered by a blocking and permeabilising solution in a humidified atmosphere. This solution is comprised of final concentrations of 0.1 V/V% Triton-X-100, 0.02 W/V% sodium azide, 5 V/V% horse serum (Sigma-Aldrich, Cat#H1270), 0.01 W/V% thimerosal and 0.3 W/V% bovine serum albumin (Sigma-Aldrich, Cat#A7284) in PBS at pH = 7.4.

For the detection of BrdU and IdU, tissue sections were treated for 1 hour with a solution of 2 M HCl in distilled water, followed by five rinses with PBS. After confirmation of complete neutralisation of the pH, the tissue sections were subjected to blocking and permeabilisation for the duration of 1 hour.

After blocking and permeabilisation, all tissues were washed three times in PBS for 10 minutes. Incubation with primary antibodies was always performed prior to EdU detection via the copper-catalysed click reaction as to not interfere with epitope recognition. Primary antibodies were applied overnight at room temperature in the blocking buffer with the omission of Triton X-100, in a humidified staining tray at the following dilutions:

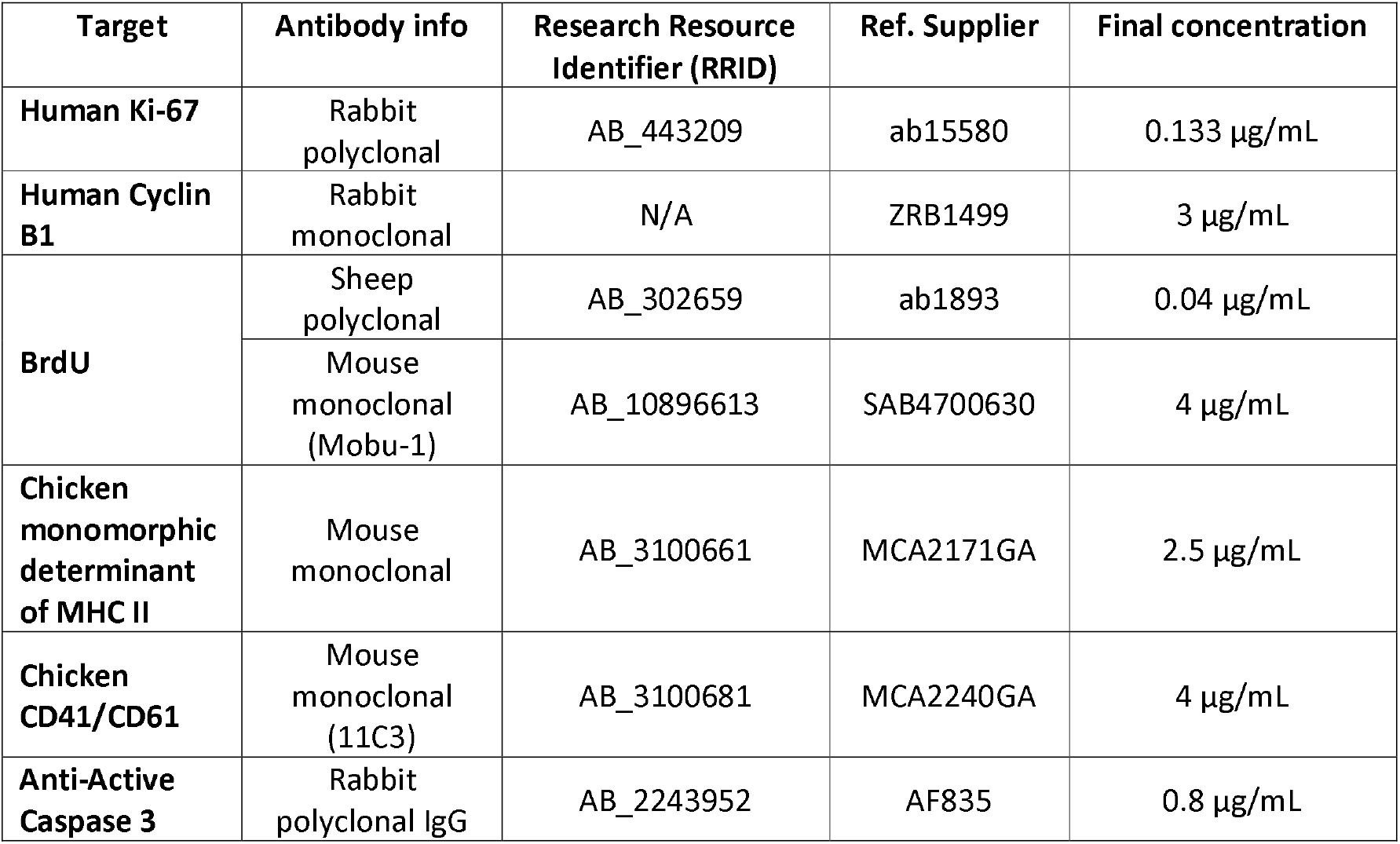

The following day, slides were washed three times in PBS and subsequently incubated with the appropriate secondary antibodies for one hour at room temperature in a humidified staining tray. All secondary antibodies were finally dissolved in the blocking buffer without Triton X-100 at a final concentration of 2 µg /mL together with Hoechst 33342 at a final concentration of 10 µg/mL

| Target species | Host species | Conjugate | Supplier | Ref. |
|---|---|---|---|---|
| **Anti-Mouse IgG** | Donkey | Cy3 | Jackson Immuno | 715-165-150 |
| **Anti-Rabbit IgG** | Donkey | Cy3 | Jackson Immuno | 711-165-152 |
| **Anti-Rabbit IgG** | Donkey | Cy5 | Jackson Immuno | 711-175-152 |
| **Anti-Chicken IgY** | Donkey | FITC | Jackson Immuno | 703-095-155 |
| **Anti-Goat IgG** | Donkey | Cy3 | Jackson Immuno | 705-165-147 |

After secondary antibody labelling tissues were washed three times in 0.1M PBS and either subjected to further copper-catalysed click reaction for the detection of EdU or mounted in a glycerol-based anti-fading mounting agent (Citifluor AF1, Electron Microscopy Sciences, Cat#17970-25).

### Copper-catalysed click reaction

In order to detect DNA-incorporated (F-ara-) EdU, permeabilised tissues were covered and incubated at room temperature with the staining buffer comprised out of a final concentration of 2.5 µM Azide fluorophore-conjugated dye (CF®647, Biotium, Cat#92084 or Alexa Fluor™ 647 Azide, ThermoFisher, Cat#A10277, 6-FAM-Picolyl-Azide, Baseclick, Cat#BCFA-247), 1 mM copper sulphate and 5 mM L-Ascorbic acid sodium salt (Thermo Scientific Chemicals) dissolved in PBS. Incubation durations of 30, 60 and 120 minutes were evaluated with no discernible differences in signal-to-noise ratio nor background staining. After the click reaction, tissues were washed three times in PBS and mounted (see above).

### Wheat-germ-agglutinin labelling

Permeabilised and blocked tissue sections were incubated for 45 minutes with Alexa Fluor^TM^ 488-conjugated WGA (Invitrogen, Cat# W11261) prior to copper-catalysed click-detection of nucleosides.

### Confocal fluorescence microscopy & image processing

Cryosections and *in vitro* cells were imaged using a Nikon Ti2 W1 spinning disk confocal microscope using the Nikon NIS-Elements software interface. Light was filtered through a quadruple dichroic mirror: 440, 521, 607, 700 nm and, for the relevant fluorophore, filtered through emission filters at 447/60 - 525/50 - 600/52 - 708/75 nm respectively. Z-stack images of tissue sections were acquired and maximum-intensity projections were made before further processing for visualisation purposes through the use of the QuPath (Version 0.5.1, available at https://qupath.github.io/) and Fiji (available at https://imagej.net/software/fiji/) software packages. Cellular classification models were built per cell line using the built-in Artificial Neural Network (ANN_MLP). Via human-in-the-loop evaluation, parameters were adjusted until the model was able to distinguish human tumour nuclei from chicken cells and tissue.

#### Identification and segmentation of nuclei within tissue sections

Detection of nuclei was performed via running the *Analyze > cell detection* function in the GUI. The image channel containing the nuclear Hoechst staining was designated as the detection channel. The following function argument values parameters were applied:

{*“requestedPixelSizeMicrons”:0.2,“backgroundRadiusMicrons”:20.0,“backgroundByReconstruction”:true,“medianRadiusMicrons”:0.0,“sigmaMicrons”:1.0,“minAreaMicrons”:20.0,“maxAreaMicrons”:250.0, “threshold”:50.0,“watershedPostProcess”:true,“cellExpansionMicrons”:0.0,“includeNuclei”:false,“sm oothBoundaries”:true,“makeMeasurements”:true}’. Nuclear sizes were estimated using the*

#### Binary thresholder

The binary classifier strategy can be applied on a pixel-wise or a nucleus-wise scale and classifies pixels or nuclei to either ‘Chicken’ or ‘Human’ based upon a fixed intensity threshold in the anti-IgY IF channel and is manually determined. In brief, pixels or nuclei whose (mean) intensity exceeds this threshold value are assigned the ‘Chicken’ class, whereas intensities lower than or equal to this threshold lead to the ‘Human’ class assignment. The functionality can be called through the *‘Classify > object classification > create single measurement classifier*’ or via ‘*Classify > pixel classification > create thresholder’*. The actual value of the intensity threshold was determined by the analysing researcher and remained constant for the complete set of images in the project. The threshold value was determined via trial-and-error visual inspection of its classifier outcomes on the project images.

#### Creating and training species-specific classifier models

For each cell line separately, one image training region (500 by 500 pixels) per microscopic field-of-view was extracted using the *‘Create Training Image’* command. These training images were subsequently manually annotated and different ROIs types were defined. First, clear tumour nodules were annotated and manually attributed the ‘Tumor’ class. In a similar fashion, non-tumour cells and tissue (e.g., extracellular space, epithelial embryonic structures) were assigned the ‘Stroma’ class which served as a placeholder for chicken-origin tissues and cells. Non-tissue areas were annotated and assigned the class ‘Other’.

For object-based classification, the function under *‘Classify > object classification > train object classifier*’ was called. The classifier model used is the artificial neural network (ANN_MLP) model. Depending on the inclusion the different colour channels, the relevant features were selected. For example, a two-channel trained classifier based upon the IgY and Ki-67 IF channels was instructed not to take into account nuclear morphological descriptors but only the intensity-derived measurements of the respective channels. For pixel-based classification, the function under *‘Classifiy > train pixel classifier’* was called. As mentioned, the classifier models were instructed to classify the objects (nuclei) or pixels in one of three classes ‘Tumor’, ‘Stroma’, or ‘Other’. Initial evaluation of the classification performance was assessed visually by applying the model to the generated training images and careful inspection of the ability to correctly segregate tumour cells from chicken cells. Further information about the specific parameters of each applied model are available upon request.

### Statistical analysis

QuPath measurement data were exported in .tsv format and imported into Graphpad Prism 10 (available at https://www.graphpad.com/) for further statistical analysis. Prior to any analysis, data distributions were assessed for normality. For normally distributed data, between-group differences in mean values for n > 2 groups were assessed via two-sided hypothesis testing using one-Way ANOVA followed by Tukey’s post-hoc test for multiple comparisons at a significance threshold α = 0.05. For n < 2 groups, Welch’s t-test was applied. Unless otherwise specified, mean values are plotted along with the individual data points. Error bars represent standard deviation.

## Author contributions

Conceptualisation: RC; Funding Acquisition: JPB, JPT; Methodology: RC, DJ; Investigation: RC, DJ; Supervision: JPT, JPB; Writing – Original Draft: RC; Visualisation: RC; Writing – Review & Editing: RC, JPT, JPB

## Acknowledgements

All experimental procedures were performed in the Laboratory of Cell Biology and Histology, University of Antwerp in accordance with the Directive 2010/63/EU of the European Parliament and the Belgian Royal Decree of 29 May 2013.

The authors would like to thank Grégoire Defrenne and Anass Moumouh for their voluntary practical assistance in the initial experiments concerning the nucleoside labelling in the CAM model.

