## Supplementary Figures for "Nucleoside analogue labelling to study the cell cycle of pancreatic cancer xenografts in the chicken embryo model"

**Supplementary Figure S1**

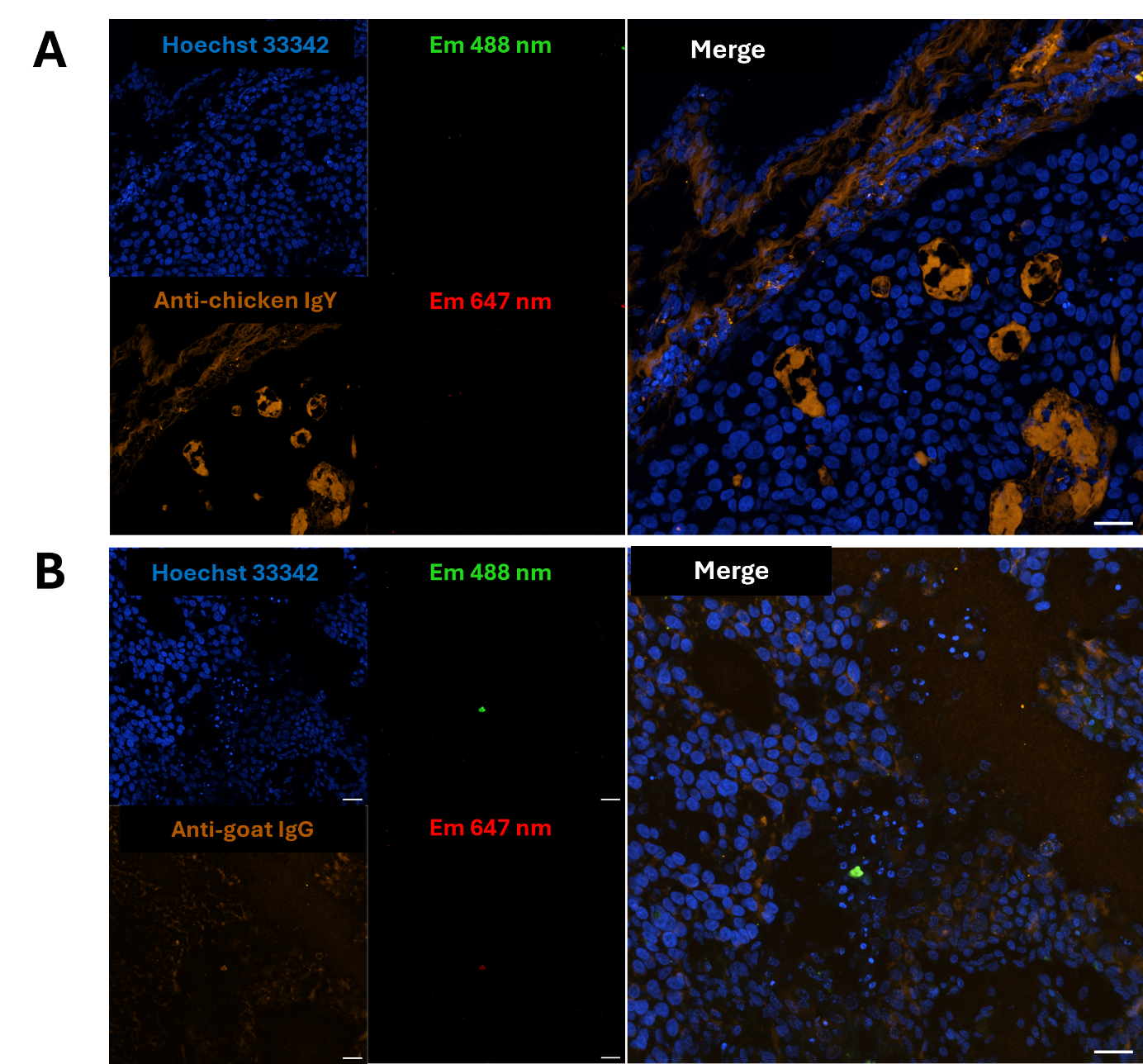

Comparison of the staining pattern of anti-chicken IgY compared to the non-specific pattern when anti-IgG secondary antibodies directed to a non-relevant species (goat in this case) are applied on BxPC-3 xenograft tissue sections. Tissues were stained with the respective Cy3-conjugated secondary (**orange**) antibodies and Hoechst 33342 (**blue**), following blocking and permeabilistation, as described. Em 488 nm (**green**) and Em 647 nm (**red**) illustrate the autofluorescence in these tissue sections when illuminated with higher-than routine illumination intensities at these excitation wavelengths. Images were acquired through a 40x water objective. Scale bars represent 25 µm.

**Supplementary Figure S2**

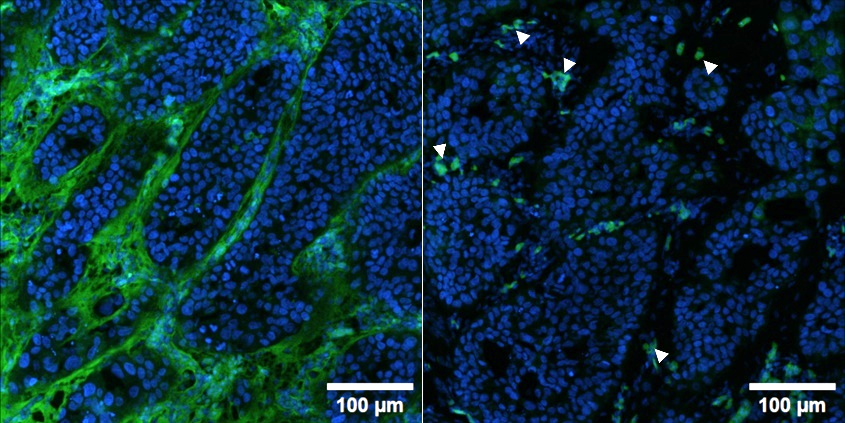

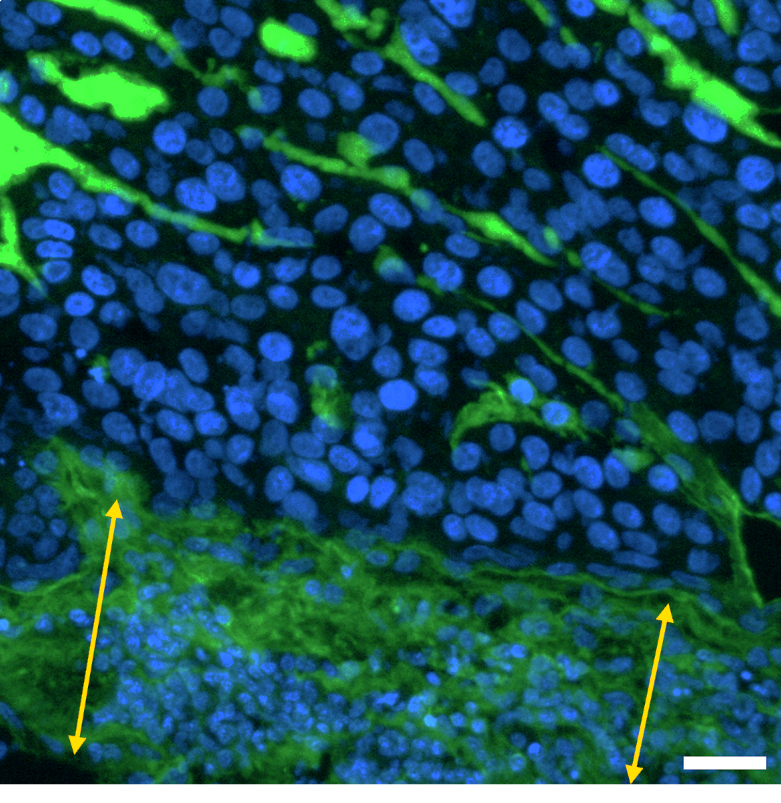

Comparison of the application of anti-IgY IF labelling (left, **green**) versus no IF labelling (middle) in BxPC-3 sections. Highly auto-fluorescent (nucleated) embryonic erythrocytes can be observed throughout the tissue (**arrowheads**) in absence of stringent permeabilisation. Anti-IgY immunofluorescence labelling (**green**) intensely stains the epithelial chicken CAM layers (**arrows**), as well as extracellular components that separate cellular tumour noduli/strands in a cryosection of a BxPC-3 xenograft at ED14. Nuclei are stained with Hoechst 33342 (**blue**). Images acquired through a 20x objective. Scale bar indicates 25 µm unless otherwise indicated.

**Supplementary Figure S3**

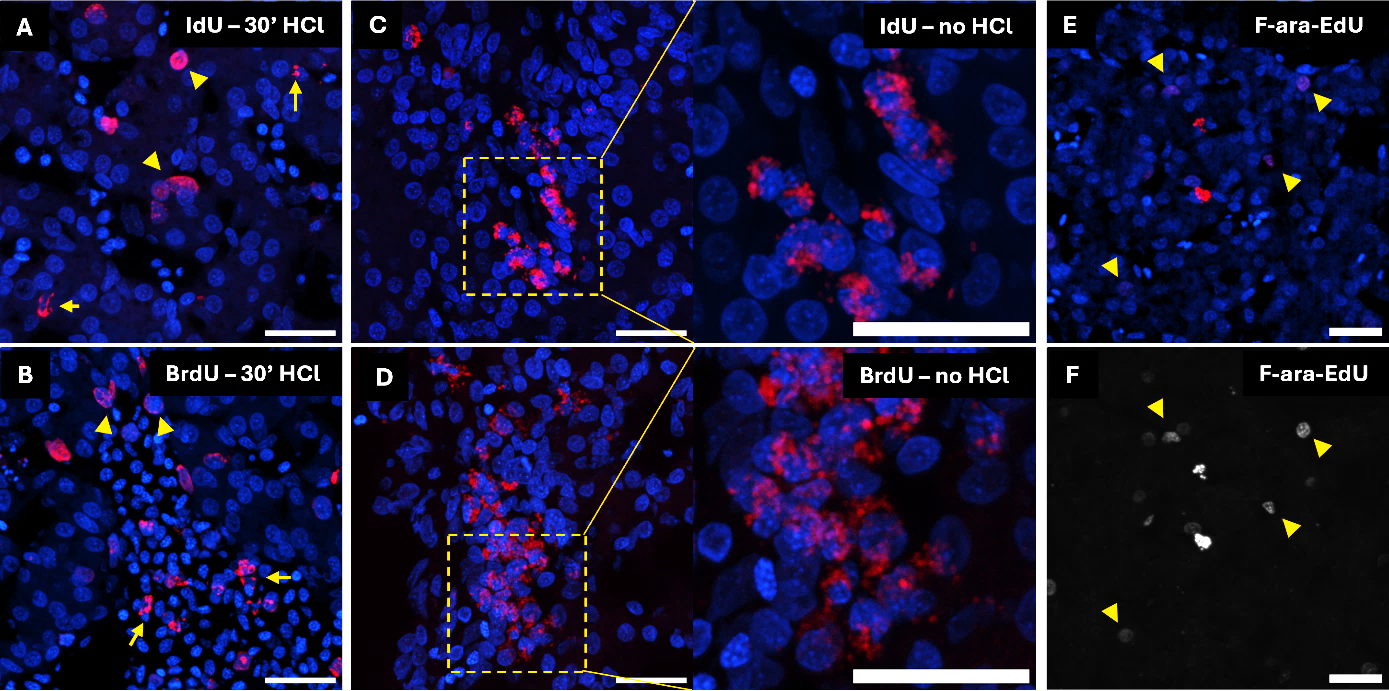

The extranuclear nucleoside signal is affected by HCl denaturation. Confocal (immuno-)FM imaging of non-grafted chicken embryo liver cryosections showing nuclear (**arrowheads)** as well as extranuclear (**arrows**) localisation of the nucleoside signal (**red**). **A-B.** IF detection of the halogenated analogues in sections denatured with 2 M HCl for 30 minutes. **C-D**. IF detection without prior denaturation. **E**. F-ara-EdU incorporation was detected in chicken embryo nuclei (**arrowheads**) and within the perinuclear environment following a 3 hour labelling period. **F.** Gray-value representation of panel **E** highlighting the low rate of nuclear F-ara-EdU incorporation. Nuclei are stained with Hoechst 33342 (**blue**). Scale bars = 25 µm. Images were acquired through a 40x objective.

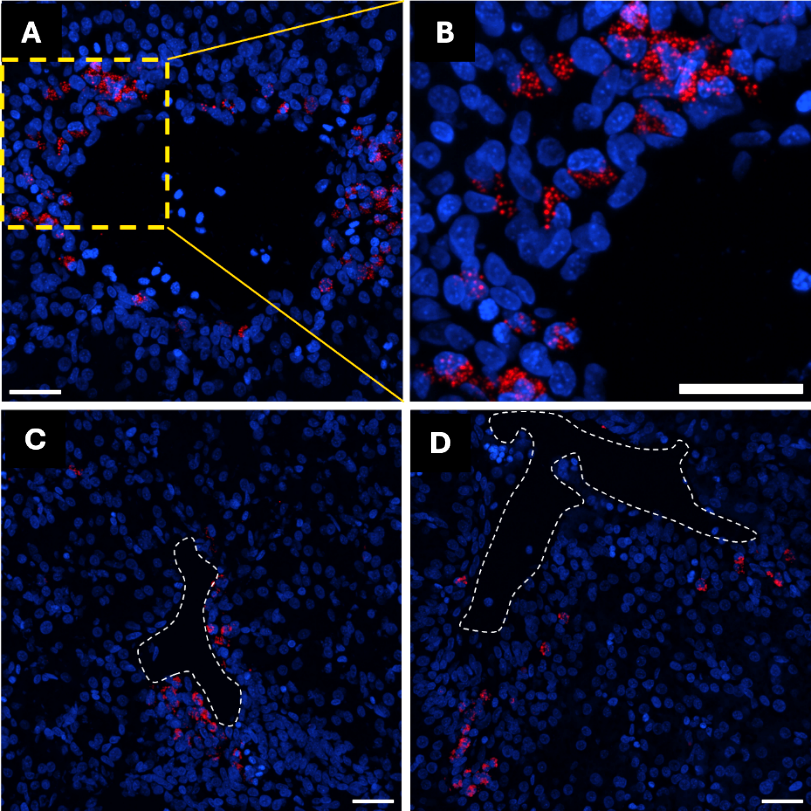
**Supplementary Figure S4**

Confocal IF imaging of non-grafted chicken embryo liver cryosections acquired at 40x magnification showing the typical localisation of cell clusters containing the nucleoside analogues (**red**) near large vessels. This was observed for F-ara-EdU (**A-B**) as well as for BrdU (**C**) and IdU (**D**) under non-denaturing conditions. Nuclei are stained with Hoechst 33342 (**blue**). Scale bars = 25 µm.

**Supplementary Figure S5**

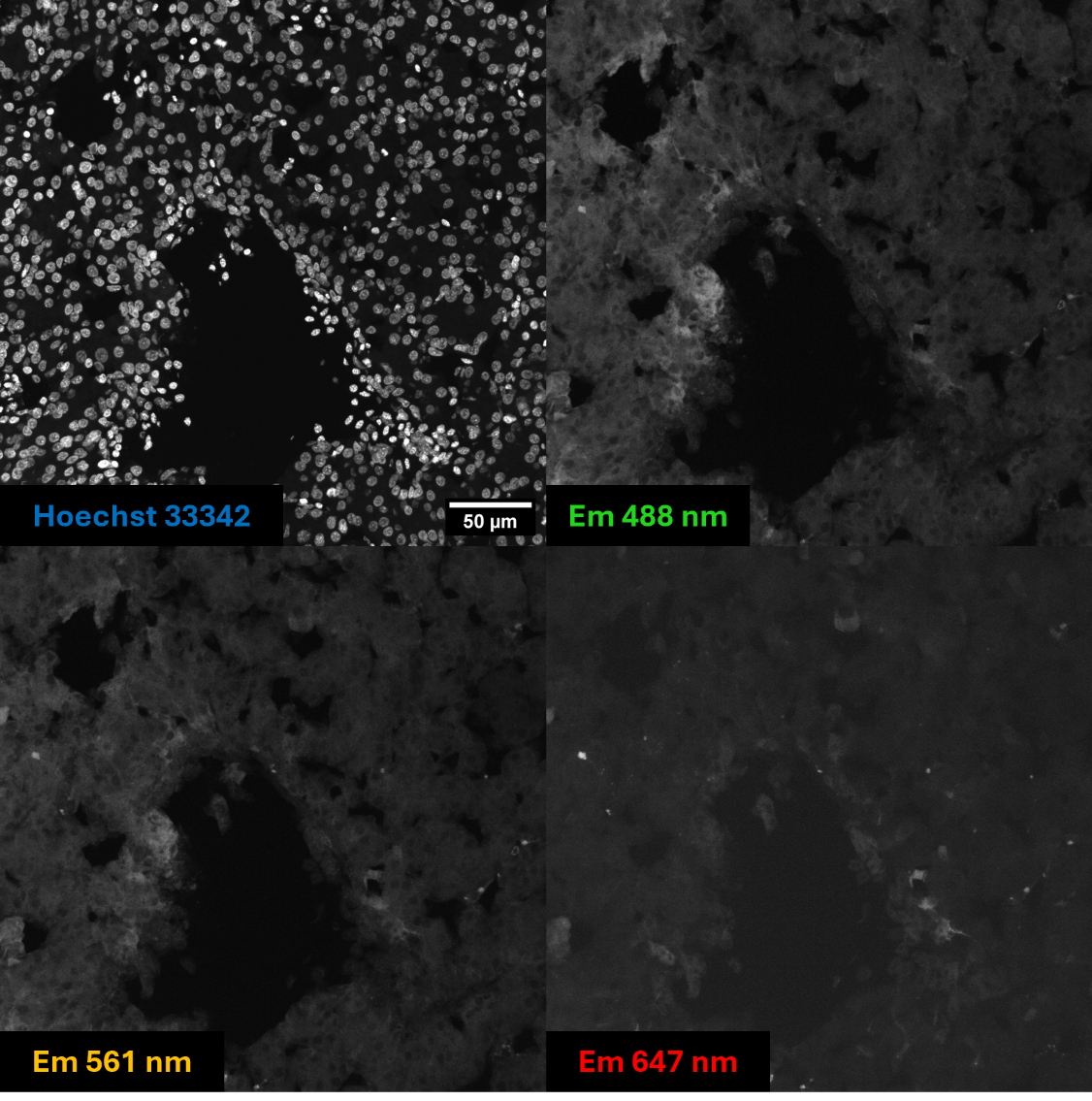

Confocal microscopic image of a non-nucleoside containing chicken liver embryo tissue section (ED15), subjected to the click-detection reaction as described. Image panels reflect the patterns of endogenous autofluorescence at each displayed emission wavelength in addition to nuclear labelling through Hoechst 33342. No discrete, granular cytoplasmic autofluorescent signals similar to those observed following nucleoside labelling are observed at all wavelengths. Images acquired through a 40x objective.

**Supplementary figure S6**

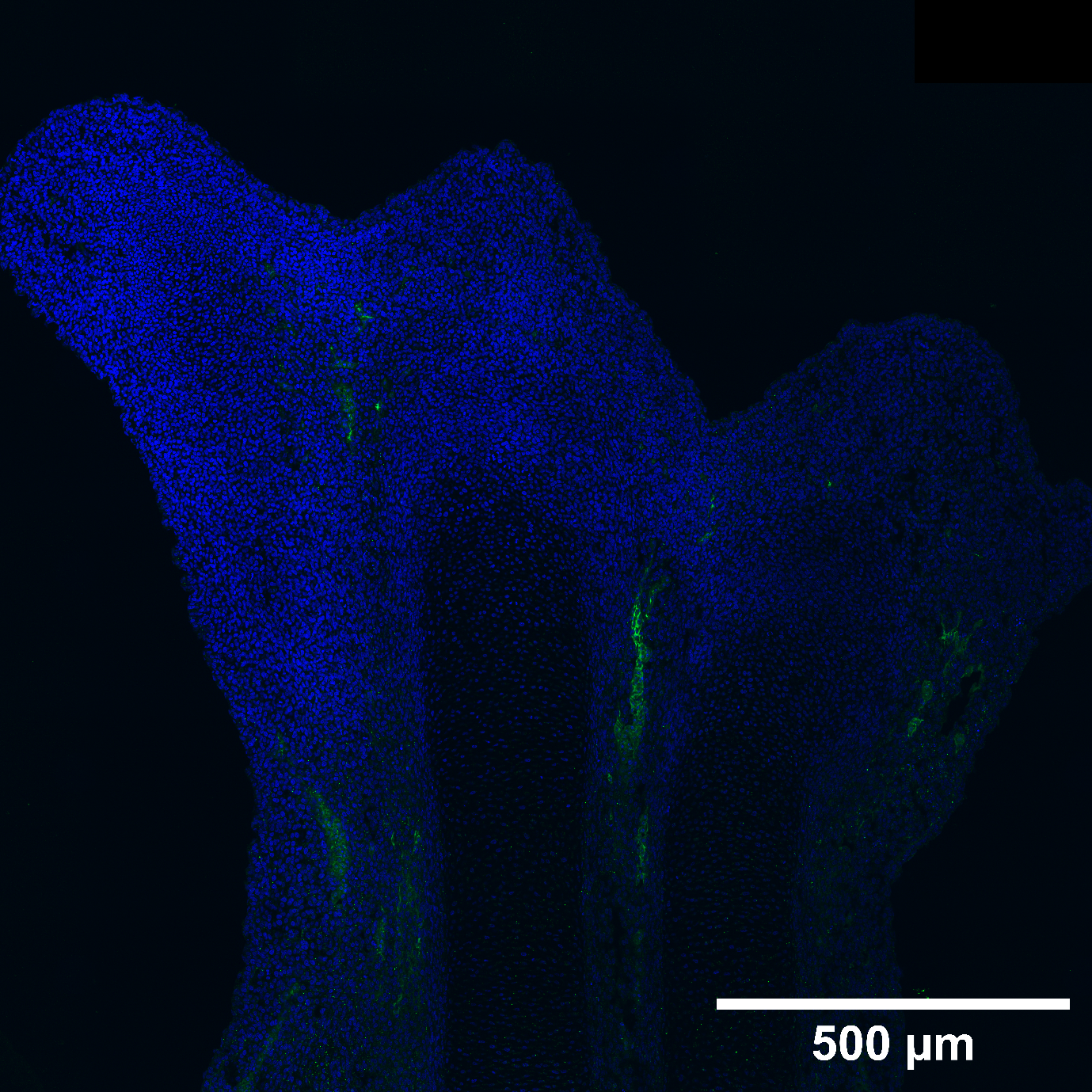

Validation of the primary anti-active caspase-3 antibody for the detection of apoptotic cells in a chicken embryo frontal limb bud (ED7; HH30-32). The active-caspase-3 signal (**green**) is localised to the interdigital apoptotic regions, confirming the antibodies’ specificity to detect apoptosis in chicken cells. The image was generated via live-stitching of multiple fields-of-view, acquired through a 40x objective. Nuclei are stained with Hoechst 33342 (**blue**).
